# Cas9-targeted Nanopore sequencing rapidly elucidates the transposition preferences and DNA methylation profiles of mobile elements in plants

**DOI:** 10.1101/2021.06.11.448052

**Authors:** Pavel Merkulov, Sofya Gvaramiya, Roman Komakhin, Murad Omarov, Maxim Dudnikov, Alina Kocheshkova, Zakhar Konstantinov, Alexander Soloviev, Gennady Karlov, Mikhail Divashuk, Ilya Kirov

## Abstract

Transposable element insertions (TEIs) are an important source of genomic innovation by contributing to plant adaptation, speciation, and the production of new varieties. The often large, complex plant genomes make identifying TEIs from short reads difficult and expensive. Moreover, rare somatic insertions that reflect mobilome dynamics are difficult to track using short reads. To address these challenges, we combined Cas9-targeted Nanopore sequencing (CANS) with the novel pipeline NanoCasTE to trace both genetically inherited and somatic TEIs in plants. We performed CANS of the *EVADÉ* (*EVD*) retrotransposon in wild-type *Arabidopsis thaliana* and rapidly obtained up to 40x sequence coverage. Analysis of hemizygous T-DNA insertion sites and genetically inherited insertions of the *EVD* transposon in the *ddm1* genome uncovered the crucial role of DNA methylation in shaping *EVD* insertion preference. We also investigated somatic transposition events of the *ONSEN* transposon family, finding that genes that are downregulated during heat stress are preferentially targeted by *ONSEN*s. Finally, we detected hypomethylation of novel somatic insertions for two *ONSEN*s. CANS and NanoCasTE are effective tools for detecting TEIs and exploring mobilome organization in plants in response to stress and in different genetic backgrounds, as well as screening T-DNA insertion mutants and transgenic plants.

## Introduction

Transposable elements (TEs) are a diverse group of genomic elements that can jump across the genome via RNA intermediates (Class I transposons, or retrotransposons) or via a “cut-and-paste” mechanism (Class II, DNA transposons). TEs have been identified in almost all living organisms, where they and their remnants can comprise up to ∼93% of the genome (Rabanus-Wallace *et al*., 2021). Long terminal repeat (LTR) retrotransposons are a major group of TEs in plants with hundreds of ancient as well as very recent insertions (Baduel *et al*., 2021). TE insertions (TEIs) can alter gene expression, creating a new transcriptional repertoire and causing genomic reorganization (Chuong *et al*., 2017). The effects of TEs on genomic, transcriptomic and proteomic activity have made them a major force in the diversity and evolution of plant species (Lisch, 2013). TEIs are classified as genic (exonic and intronic) or intergenic (Sultana *et al*., 2017) based on their locations in the genome. TEIs in or near genes are often more deleterious than single nucleotide polymorphisms (SNPs) and may create null alleles. However, the true consequences of TEIs on gene function are variable, ranging from complete inactivation to the generation of novel transcriptional programs.

Progress in pangenome sequencing and genome assembly algorithms has helped unravel the genome-wide TEI landscape in natural plant populations and germplasm (Domínguez *et al*., 2020; Baduel *et al*., 2021). These studies demonstrated that TEIs are an important source of novelty for plant adaptation (Domínguez *et al*., 2020). For example, recurrent insertions of TEs in *FLOWERING LOCUS C* (*FLC*; a key determinant of flowering time in plants) in different Arabidopsis (*Arabidopsis thaliana*) accessions have contributed to the local adaptation of this species (Baduel *et al*., 2021). A recent genome-wide association study (GWAS) using 602 resequenced genomes of wild and cultivated tomatoes (*Solanum lycopersicum*) revealed that at least five of 17 analyzed traits were associated with polymorphisms caused by TEIs (Domínguez *et al*., 2020), including fruit color and leaf morphology. Hence, ongoing TE activity has also been widely exploited by plant breeders, as many novel and low-frequency alleles of the genes involved in agronomically important traits originated via TEIs (Kobayashi *et al*., 2004; Jiang *et al*., 2009; Butelli *et al*., 2012; Domínguez *et al*., 2020).

TE transposition activity in plant cells is under strict epigenetic control, and most TEs are silenced under non-stress conditions (Slotkin and Martienssen, 2007; Nuthikattu *et al*., 2013). To overcome this limitation for TE investigation, mutants that are defective in key genes of TE epigenetic control systems have been obtained. These plants are fertile and have elevated TE activity, making them indispensable tools for studying TE biology (Mirouze *et al*., 2009; Tsukahara *et al*., 2009; Panda and Slotkin, 2020).

Identifying TEIs in the genome is challenging, primarily because most plant genomes possess a large amount of repetitive DNA carrying many of these elements. The detailed picture of the organization of this ‘dark side’ of the genome is difficult to explore using a short-read approach. Long-read sequencing using PacBio and Nanopore technologies provides a dramatic increase in the resolution of the structures of repetitive DNA sequences (Jung *et al*., 2019; Kirov *et al*., 2021*a*). While whole-genome sequencing (WGS) theoretically allows all TEIs in an individual plant genome to be identified, its application in plants with large genomes is still quite expensive. WGS is even more difficult when TEIs in a plant population must be identified (Handsaker *et al*., 2011). Therefore, several approaches have been proposed to capture specific genomic loci possessing TEIs without the need to sequence the rest of the genome (Li *et al*., 2019). Enriched DNA samples are usually sequenced using short reads. Therefore, identifying TEIs that have occurred in repeat-rich regions such as centromeres and in other TEs is challenging. Enrichment for a specific sequence is mostly based on oligo-probe-mediated ‘fishing’ of target DNA fragments (Williams-Carrier *et al*., 2010; Quadrana *et al*., 2016). A recently developed microfluidics-based method (Xdrop) was shown to be effective for capturing the integration sites of human papillomavirus 18 (Madsen *et al*., 2020). While these approaches are effective, they require either a PCR amplification step (which may introduce chimeric reads) or the use of specific equipment (for the Xdrop method).

The combination of TE capture and Nanopore sequencing has greatly improved integration site mapping (Li *et al*., 2019; Madsen *et al*., 2020). A novel enrichment approach combining Cas9-mediated adaptor ligation and long-read Oxford Nanopore sequencing (CANS) was recently proposed (Gabrieli *et al*., 2018; Gilpatrick *et al*., 2020; Stangl *et al*., 2020). This method is based on the finding that Nanopore sequencing adaptors preferentially ligate to Cas9-cleaved DNA sites of artificially dephosphorylated genomic DNA. The advantages of this approach over oligo-based enrichment include the lack of a PCR amplification step, the ability to detect DNA methylation and the shorter time needed to complete the protocol. CANS has been used to sequence target genes in humans (Gilpatrick *et al*., 2020) and plants (López-Girona *et al*., 2020; Kirov *et al*., 2021*b*). The use of CANS to identify TE integration sites in human cells was recently reported (McDonald *et al*., 2021). However, the use of CANS for identifying TEIs in plants has not yet been reported.

Here we used CANS and developed a novel pipeline (NanoCasTE) to identify T-DNA and somatic TEIs, as well as genetically inherited TEIs, in Arabidopsis. Using the Arabidopsis *ddm1* (*decrease in DNA methylation 1*) mutant, which has increased TE activity, we successfully detected all insertions of *EVADÉ* (*EVD*) retrotransposons in a single plant with only 0.2x genome-wide coverage by CANS reads. This coverage is an order of magnitude below that required for TEI identification based on WGS. A comparison of the distributions of hundreds of *EVD* insertions in *ddm1* (detected by CANS) and natural Arabidopsis ecotypes (Baduel *et al*., 2021) revealed a key role for methylation in shaping TEI distribution in the genome. Using CANS to detect *ONSEN* somatic insertions in wild-type Arabidopsis plants after heat stress, we demonstrated its utility for monitoring the response of the mobilome to stress without the need to obtain seeds from stressed plants. Overall, our results demonstrate that CANS and NanoCasTE are powerful tools to capture TEIs in plants, providing a way to gain a deeper understanding of TE integration landscapes in plant genomes and to explore the potential of TE mutagenesis in plant breeding.

## Results

### Cas9-mediated targeted sequencing of three copies of the *EVD* retrotransposon in Arabidopsis

To test whether CANS is suitable for tracing individual transposon insertions in plants, we chose the *EVD5* (5,333 bp) LTR retrotransposon (Mirouze *et al*., 2009), which can generate new insertions in some Arabidopsis mutants (e.g. *met1* [defective in *METHYLTRANSFERASE 1*] and *ddm1*) (Tsukahara *et al*., 2009)). *EVD5* belongs to the *EVADÉ* subfamily (*AtCOPIA93* family) of *Ty1*/*Copia* retrotransposons, which includes other *EVD* elements whose activity was not detected. Of these, *EVD1* (AT1TE41575, 1,548 bp) and *EVD2* (AT1TE41580, 5,336 bp) are densely located elements with high sequence identity (∼99%) to *EVD5* (Mirouze *et al*., 2009).

The CANS workflow consists of several steps: (1) isolation and size selection of high amounts (5–10 µg) of high-molecular weight genomic DNA; (2) design and synthesis of single guide RNA (sgRNA) by *in vitro* transcription; (3) Cas9-sgRNA complex assembly; (4) genomic DNA cleavage by Cas9-sgRNA complexes and dA-tailing; (4) Nanopore adaptor ligation and (5) Nanopore sequencing. The detailed protocol of this procedure is available on protocols.io (https://www.protocols.io/private/DE1CFDE8C8FF11EBA7DA0A58A9FEAC02).

For the CANS experiments, we designed a set of seven sgRNAs located in LTRs and inter-LTR regions of *EVD5* and *EVD2*. We predicted that one of these sgRNAs also recognizes the *EVD1* copy. Because CANS is strand-specific, we designed sgRNAs to direct Nanopore sequencing to sequence (1) *EVD* retrotransposons *per se* and (2) upstream and downstream sequences flanking retrotransposon insertions. We transcribed the sgRNAs *in vitro* as single-guide RNAs (sgRNAs). We carried out two runs on a MinION sequencer using seven and five sgRNA sets, respectively. The pilot experiment using all sgRNAs as one mixture yielded 15,777 reads (144 Mb total bases), with 231 (1.4%) on-target reads detected after 4.4 h of sequencing. Because many short reads were produced from the internal sgRNA cutting site, for our second experiment, we used five of the most efficient sgRNAs divided into two mixtures to prevent the generation of short reads. These sgRNAs, designated sgRNA1 to sgRNA5, comprised one LTR sgRNA (sgRNA1, sgRNA mixture 1) and four sgRNAs (sgRNA mixture 2) located in the inter-LTR region (Figure 1).

**Figure 1.**
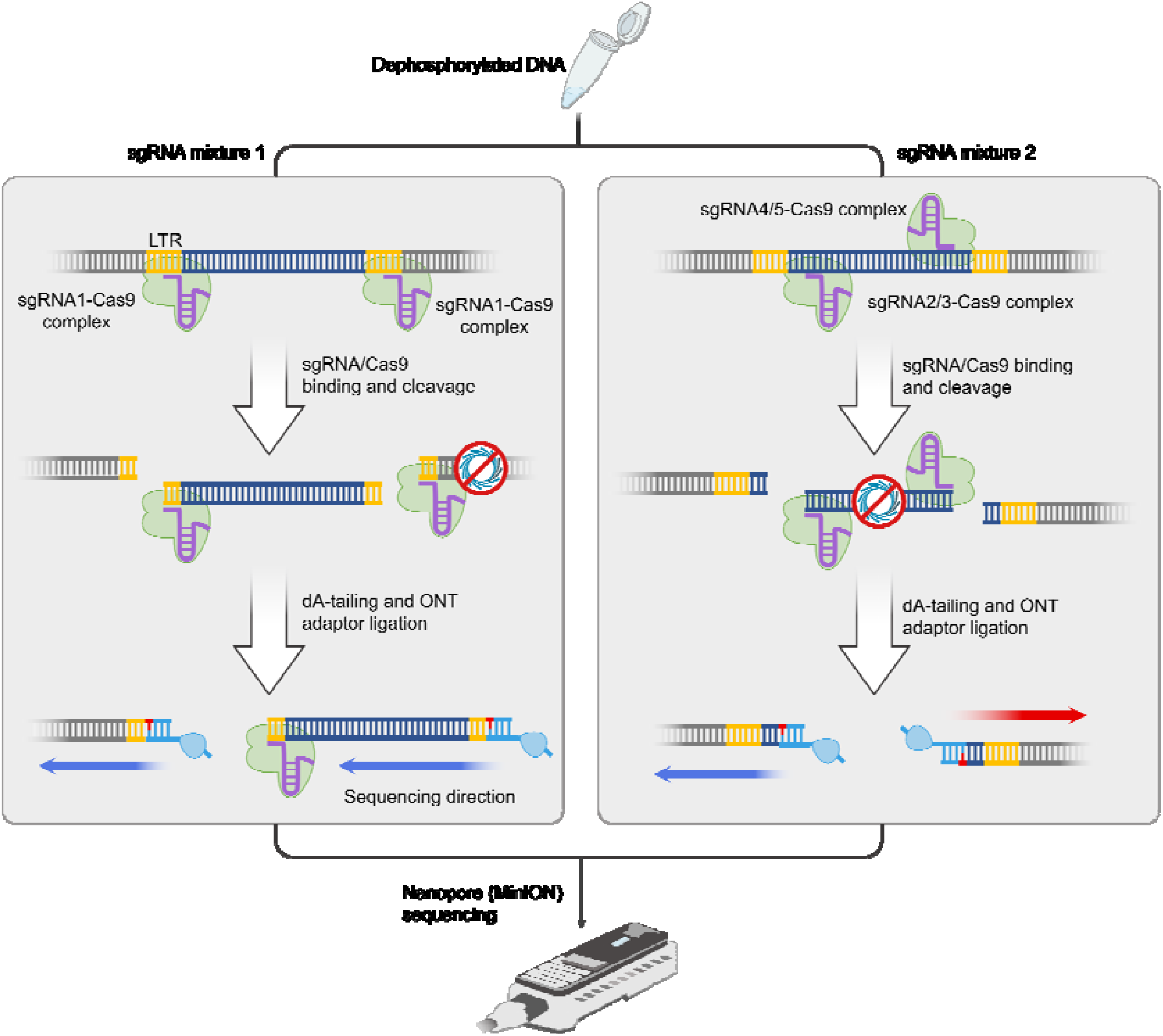
Schematic diagram depicting the identification of CANS of LTR retrotransposon insertions using five sgRNAs divided into two mixtures. Blue and red arrows indicate the strand specificity (negative and positive strand, respectively) of sequencing depending on sgRNA orientation.

After 4 hours of sequencing on the MinION flow cell the experiment using five sgRNAs yielded 88,000 reads (207 Mb total bases) and 259 (0.3%) of them were on-target reads. Both CANS runs using Arabidopsis Col-0 genomic DNA resulted in up to 40× coverage of the target retrotransposons *EVD5*, *EVD1* and *EVD2* (Figure 2A, 2B). Mapped reads from the first and second runs covered regions of 35,000 bp and 14,000 bp, respectively, including target *EVD*s and their flanking regions. The broad coverage of the flanking sequences demonstrates the advantage of CANS for detecting TEIs in the genome. Inspection of the sgRNA locations and read distribution revealed that CANS has high strand specificity, with up to 82% of reads mapped to strands with 3’ ends unprotected by Cas9 and possessing PAM (NGG) sequences. This feature is another advantage of CANS, as it allows the coverage of the target region to be increased. We also noticed that the bulk of reads from internal parts of *EVD2* and *EVD5* were unambiguously assigned to the corresponding *EVD* copy, corroborating previous reports that cDNA Oxford Nanopore (ONP) reads have high mappability (Panda and Slotkin, 2020).

**Figure 2.**
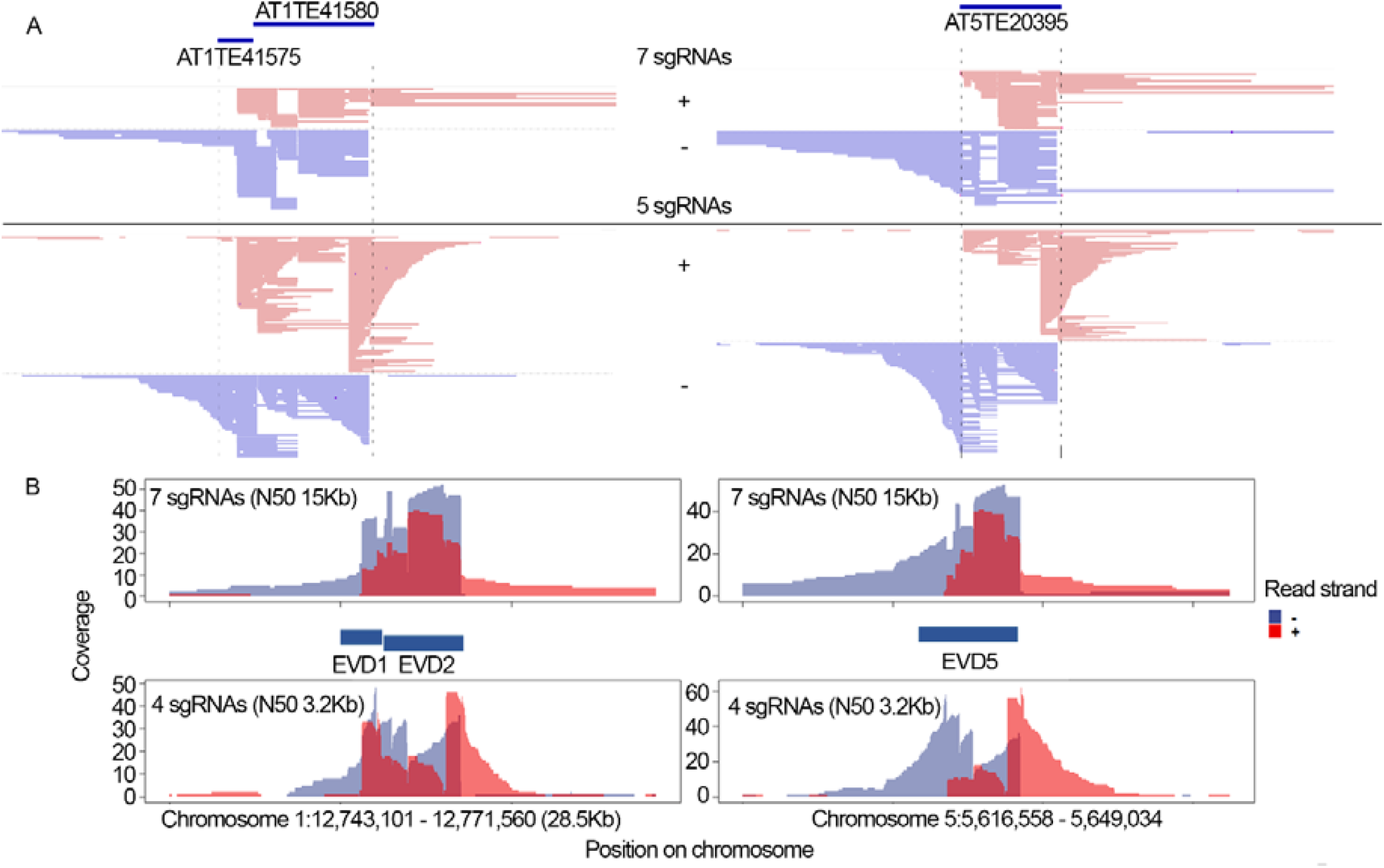
Targeted enrichment of *EVD* loci following CANS using a set of 7 or 5 sgRNAs. (**A**) Integrative Genomics Viewer (IGV) window of Nanopore reads aligning to *EVD1* (AT1TE41575), *EVD2* (AT1TE41580) and *EVD5* (AT5TE20395) retrotransposon copies. (**B**) Coverage plots are shown for *EVD1*, *EVD2* and *EVD5* using forward (red) and reverse (blue) reads obtained following CANS with 7 and 5 sgRNAs.

Thus, using CANS, we sequenced *EVD1*, *EVD2* and *EVD5* retrotransposon copies and their flanking sequencing with up to 40× coverage, demonstrating the advantage of this approach for target TE sequencing.

### Detection of T-DNA and new *EVD* insertions in the genome using CANS and a new pipeline

To assess whether non-reference *EVD* insertions could be detected using CANS, we exploited the Arabidopsis *ddm1* mutant, whose *EVD* activity was previously described (Tsukahara *et al*., 2009). We created transgenic T_2_ *ddm1* plants carrying T-DNA insertions containing the *EVD5 GAG* open reading frame (*G-ddm1-3* plants). The T-DNA is recognized by one of the five *EVD* sgRNAs, allowing us to simultaneously detect *EVD* insertions and T-DNA integration sites and to obtain information about the sufficiency of CANS sequencing depth. We carried out CANS using five sgRNAs and generated 4,420 reads with an N50 ∼11 kb. Following read mapping to the Col-0 reference genome, we examined the original *EVD* copy on chromosome 5 to estimate the sequence coverage (Figure 3A). The *EVD* sequence reached up to 400× coverage with 1,170 mapped reads (25% of all CANS reads) (Figure S1). Notably, the reads spanning *EVD* sequences from LTR to LTR could not be assigned to a certain novel *EVD* insertion, as all *EVD* copies had identical sequences. Consequently, the observed *EVD* coverage primarily resulted from the alignment of Nanopore reads that originated from different *EVD* insertions in the genome.

**Figure 3.**
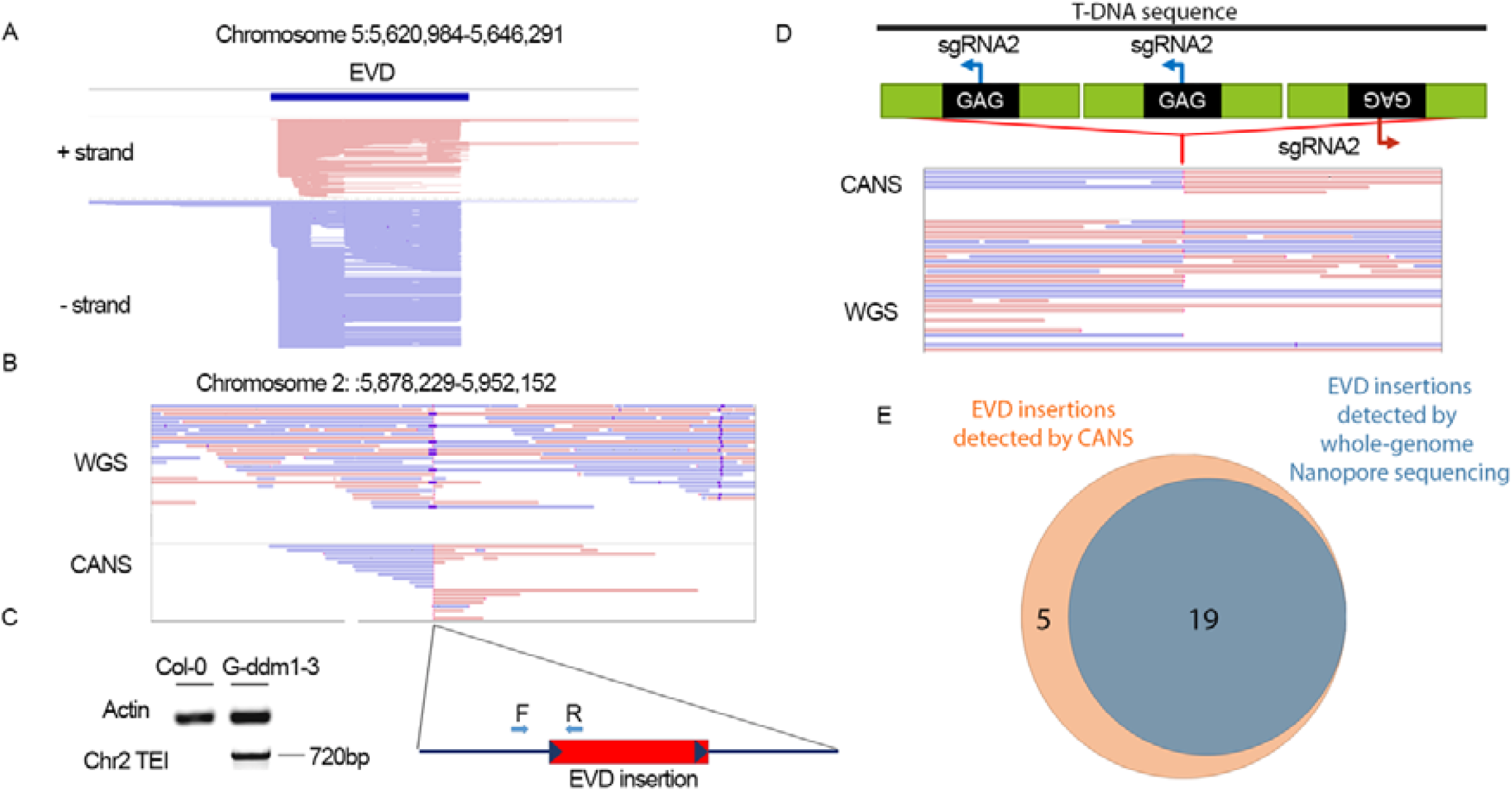
*EVD* coverage and detection of insertions in a G-*ddm1-3* plant using CANS. (**A**) IGV snapshot of coverage of the original copy of *EVD* on chromosome 5 using forward (red) and reverse (blue) ONT reads obtained after CANS of a G-*ddm1-3* plant. (**B**) IGV browser snapshot of one of the 25 genetically inherited *EVD* insertions detected by whole-genome Nanopore sequencing and CANS. (**C**) PCR validation of the *EVD* insertion on chromosome 2 (left) using primers based on the *EVD* and TEI flanking region (right). (**D**) Structure of the inserted T-DNA (upper panel), sgRNA2 positions and its coverage based on CANS and whole-genome Nanopore sequencing reads. (**E**) Venn diagram of the number of *EVD* insertions detected in a G-*ddm1-3* plant by CANS and whole-genome Nanopore sequencing, followed by analysis using the nanotei pipeline.

To automate the detection of TE and T-DNA insertions, we developed a novel pipeline called NanoCasTE (see Materials and Methods). By applying NanoCasTE to the CANS dataset, we detected 24 *EVD* insertions. Of these, 19 were high-confidence TEIs (supported by two or more reads). An example of one TEI located on chromosome 2 that was homozygous according to the WGS data is shown in Figure 3B. We validated the existence and position of this TEI by PCR with primers designed based on *EVD* and TEI flanking sequences (Figure 3C). To support the notion that the coverage we obtained was sufficient for *EVD* TEI identification, we analyzed *EVD*-genome junctions in the sequencing data we obtained and determined that they had ∼8× coverage, which is sufficient for insertion detection (Figure 3A).

We investigated whether T-DNA insertions sites could be detected by CANS. The T-DNA sequence (*EVD5 GAG*) possesses the recognition site for a single sgRNA (sgRNA2, Figure 3D). Using CANS, we identified one T-DNA insertion site represented with 4x coverage (Figure 3D). We confirmed the presence of this T-DNA insertion site by whole-genome Nanopore sequencing followed by running the nanotei pipeline (Kirov *et al*., 2021*a*), which also revealed that the T-DNA insertion was hemizygous. These results demonstrate that even using a single sgRNA, a hemizygous T-DNA integration site in the Arabidopsis genome can be detected by CANS.

Next, we estimated the efficiency of CANS. To this end, we performed whole-genome Nanopore sequencing of the same G-*ddm1-3* plant that was used for CANS, followed by TEI calling using the nanotei pipeline. We identified 19 *EVD5* insertions supported by 2 or more WGS Nanopore reads. A comparison of TEIs identified by CANS and WGS Nanopore sequencing showed that all 19 TEIs identified by WGS are among the high-confidence TEIs captured by CANS (Figure 3E). These results indicate that even the low genome coverage (0.2× in our experiment) of CANS may be sufficient to capture all T-DNA and target transposon insertions in Arabidopsis, highlighting the good sensitivity of this method. Additionally, the newly developed NanoCasTE pipeline enables the rapid identification of novel TEIs from CANS data.

### Capture of somatic insertions in a *ddm1* population reveals differences in chromosome distribution compared to wild-type accessions

Some TEs are thought to be preferentially inserted into certain genomic regions, but the current picture of TEI distribution has been shaped by purifying selection. As CANS can detect both somatic and genetically inherited TEIs, we were interested in obtaining a picture of the *EVD* insertion landscape in real time. Accordingly, we performed CANS sequencing of *EVD* insertions in a population of approximately 50 *ddm1* plants. After 2.5 hours of MinION sequencing, we generated 18,246 Nanopore reads with N50 values of ∼5.5 kb. Using the NanoCasTE pipeline, we detected 851 TEIs, including 29 high-confidence TEIs (supported by two or more reads), in the *ddm1* population (Supplemental Data S4). We validated 13 TEIs by PCR using genomic DNA from individual *ddm1* plants, resulting in 1 to 6 plants carrying distinct TEIs. The presence of the same insertion in several plants is expected, as these 50 plants were all siblings of the same *ddm1* plant (generation F8). Analysis of the chromosome-wide distribution of the TEIs showed that they tended to clustered in the pericentromeric regions of all chromosomes, whereas the density of TEIs was significantly lower in chromosome arms (Figure 4A).

**Figure 4.**
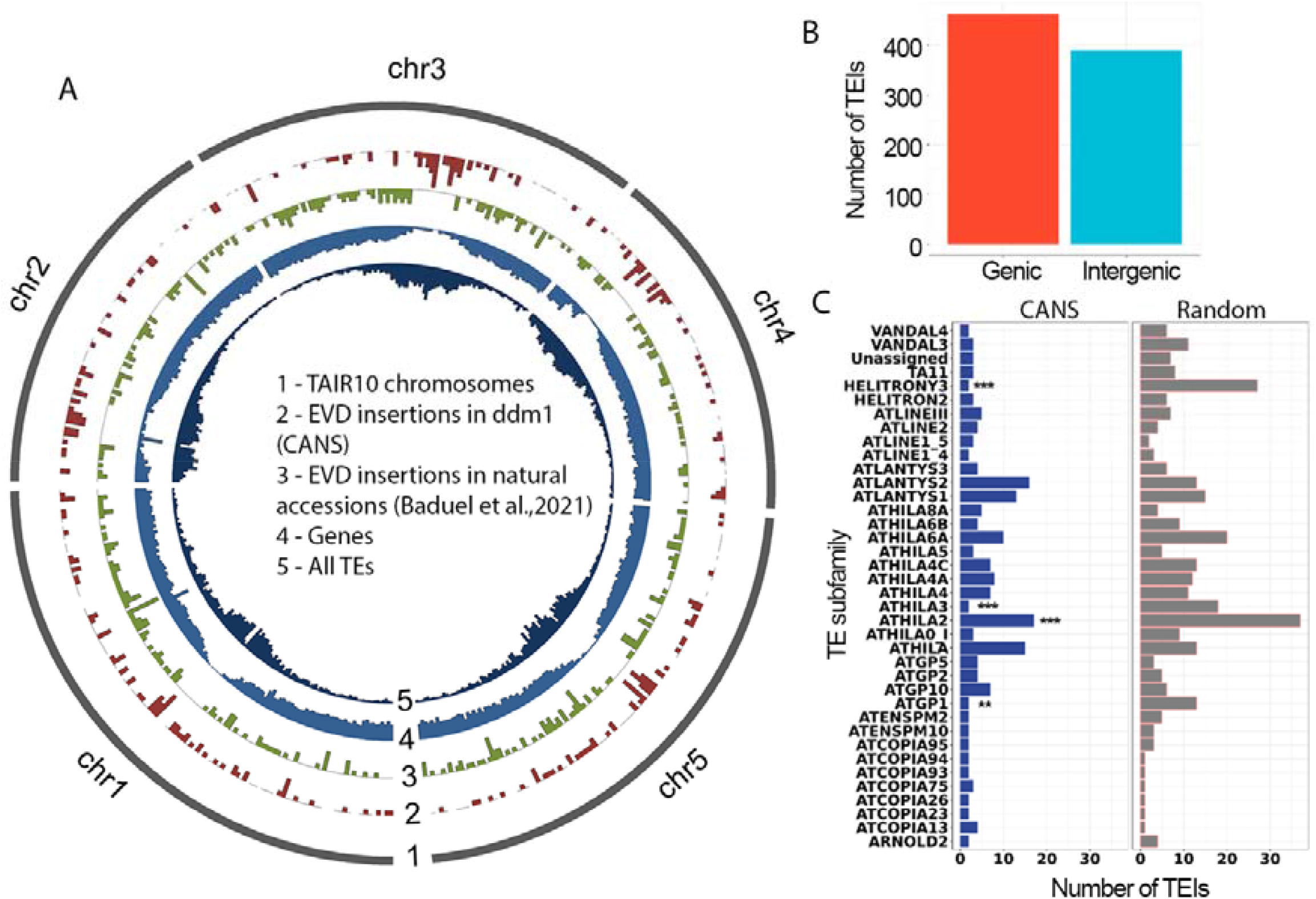
Genome organization of *EVD* insertions detected by CANS in a pooled (∼50 plants) *ddm1* sample. (**A**) Chromosome distribution of *EVD* insertions detected by CANS in the *ddm1* genome (2) and in natural Arabidopsis accessions (Baduel *et al*., 2021) (3), TAIR10 annotated genes (4) and TEs (Panda and Slotkin, 2020) (5). (**B**) Number of genic and intergenic TEIs detected by CANS. (**C**) Number of *EVD* insertions in members of different TE subfamilies in Arabidopsis based on CANS data (left panel) and random expectation (right panel).

Classification of these TEIs revealed that 54% (462) are located in genic regions, comprising 403 (87%) exonic and 59 (13%) intronic TEIs (Figure 5B). This value is expected by chance based on the total length of the genic regions in the assembled (119 Mb, TAIR10) Arabidopsis genome (68.3 M, or 57% of the sequenced genome). Gene Ontology (GO) analysis revealed no enrichment of any GO categories in the set of genes with TEIs, suggesting that most *EVD* insertions in *ddm1* are randomly distributed among genes with different functions.

**Figure 5.**
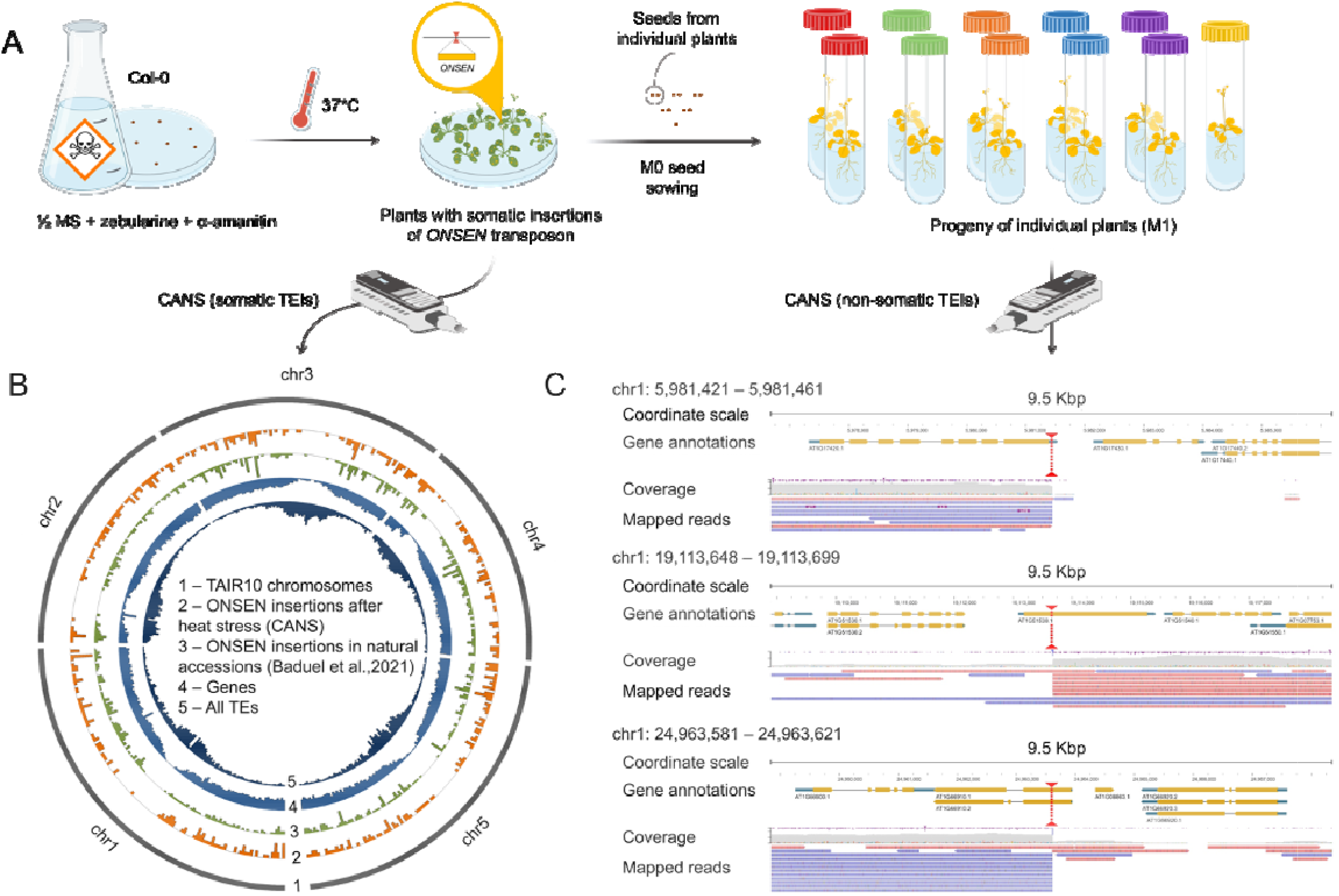
CANS detection of somatic and genetically inherited *ONSEN* insertions after heat-stress activation. **(A)** Schematic overview of the experiment examining the heat-induced activation of *ONSEN*s in the presence of zebularine and α-amanitin in wild-type Arabidopsis. (**B**) Distribution across the chromosomes of (1) *ONSEN* insertions detected by CANS in plants 48 h after heat stress, *ONSEN* insertions in natural Arabidopsis accessions (Baduel *et al*., 2021) (3), TAIR10 annotated genes (4) and TEs (Panda and Slotkin, 2020) (5). **(C)** IGV snapshots of three non-somatic *ONSEN* insertions in M1 plants detected by CANS.

Among intergenic TEIs (389, 46%), 241 TEIs (28%) were located in the sequences of other TEs. This value is significantly higher (Chi-squared test *P*-value = 0.0004) than expected by chance based on the total length of the annotated TEs in the Arabidopsis genome (24.9 Mb, or 21% (Panda and Slotkin, 2020)). Moreover, 20 TEs contained 2 or 3 different TEIs. We analyzed the number of TEIs in TEs of different subfamilies and observed that members of four subfamilies are significantly underrepresented (Fisher’s exact test *P*-value <0.01) in a set of TEs with detected TEIs: *HELITRONY3* (*P*-value = 6.78e-07), *Athila2* (*P*-value = 0.004), *Athila3* (*P*-value = 0.0002) and *ATGP1* (*P*-value=0.004) (Figure 4C). This finding can be partly attributed to the differences in chromosome distribution between TEs of these families and *EVD* TEIs. For example, most (1104 of 1399, 79%) *HELITRONY3* family members are located in chromosome arms, where the *EVD* TEI frequency is much lower than in the pericentromeric regions. *EVD* insertion preferences for other TEs cannot be explained by their chromosome location biases. In addition, we detected no DNA motif enrichment in the regions (TEI ± 500 bp) of *EVD* insertions. Therefore, we reasoned that *EVD* insertions may occur in genomic regions with a certain chromatin state, as previously shown for another transposon, *ONSEN* (Roquis *et al*., 2021).

As chromatin states are correlated with transcription status, we examined whether *EVD* insertions occur more frequently in poorly transcribed regions by performing direct Nanopore RNA sequencing of *ddm1* seedlings. Indeed, ∼90% (764) of *EVD* insertions were located in regions with low expression (<2 Nanopore reads) in the *ddm1* genome, which is significantly higher than expected by chance (Fisher’s exact test *P*-value 4.25e-18). If the chromatin state influences the insertion preferences of *EVD*s, we reasoned that the distribution of *EVD* insertions may differ between the *ddm1* mutant and wild-type genomes. To investigate this possibility, we compared the distribution of *EVD* elements between *ddm1* (851 insertions) and various wild-type Arabidopsis accessions (426 insertions of the *ATCOPIA93* family) (Baduel *et al*., 2021). Interestingly, we detected significant differences in *EVD* insertion frequency in the pericentromeric regions, with almost no insertions found in these regions in natural accessions (Figure 4A). We detected a similar distribution of *AtCOPIA93* insertions in epigenetic recombinant inbred lines (epiRILs) by Quadrana *et al*. (Quadrana *et al*., 2019). These results provide evidence that in the native genomic environment, *EVD* tends to escape pericentromeric regions, while the *ddm1* mutation skews the insertion distribution toward proximal chromosomal regions. However, the apparent depletion of *EVD* TEIs in the pericentromeric regions according to (Baduel *et al*., 2021) and (Quadrana *et al*., 2019) may be biased due to the lower mappability of short Illumina reads in repeat-rich pericentromeric regions.

Thus, using CANS coupled with the NanoCasTE pipeline, we built a high-density physical map of *EVD* insertion preferences in Arabidopsis and showed that *EVD* insertions are indeed biased toward pericentromeric regions in the *ddm1* genome. However, the distribution of *EVD* insertions strongly differed between *ddm1* and wild-type Arabidopsis accessions and the epiRILs, suggesting that DNA methylation plays a pivotal role in shaping *EVD* insertion preferences.

### Rapid detection of short-term responses of target TEs to stress

We noticed that CANS allowed both somatic and genetically inherited TEIs to be detected. This capacity could be useful for elucidating short-term responses of the mobilome to stress. To test this idea, we exploited the *ONSEN* transposon, whose transposition activity in wild-type Arabidopsis can be triggered by heat stress (Ito *et al*., 2011). To increase *ONSEN* activity and induce transgenerational inheritance, we applied heat-stress together with α-amanitin and zebularine (ZA) treatment (Figure 5A), two drugs that induce epigenetic stress. We designed six sgRNAs targeting all eight known copies of *ONSEN* (*ONSEN1*–*8*) and performed CANS experiments as described above. As an additional control, we included *EVD* sgRNAs in the same run (multiplexing). After two MinION runs on the same flow cell, we collected 175,001 high-quality reads, with 3,924 reads aligned to the *ONSEN* sequences (Figure S2).

Data analysis using the NanoCasTE pipeline revealed only one somatic *EVD* insertion in ZA-treated Col-0 plants after heat stress, which is significantly smaller than the number of *EVD* insertions in *ddm1* detected by CANS (851 TEIs, see above). These differences might be explained by a smaller coverage achieved for *EVD* in the current experiment. To test this idea, we inspected the *EVD5*-genome junctions to assess the coverage of the target region. However, we detected 60x coverage, which is 7.5 times higher than the *EVD5* coverage by reads obtained by CANS sequencing of *ddm1* plants. Therefore, the finding that only one TEI of *EVD* was detected in ZA-treated Col-0 after heat stress can be explained by the low transposition activity of this element rather than by low read yield of CANS.

We analyzed *ONSEN* insertions and identified 519 insertions (Supplemental Data S5), including one insertion identified previously as a result of a TAIR10 genome assembly error (Kirov *et al*., 2021*a*). The distribution of *ONSEN* insertions showed strong bias toward chromosome arms, while the insertion frequency in the centromere regions was very low (Figure 5B). We obtained similar results by plotting *ONSEN* insertions detected in natural Arabidopsis accessions (Baduel *et al*., 2021). Most *ONSEN* insertions (83%, 430 of 519) were located in genes, corroborating the recent finding that >90% of *ONSEN* insertions are genic (Roquis *et al*., 2021). GO enrichment analysis of the genes possessing *ONSEN* insertions (426 genes) revealed several overrepresented GO terms, including aspartyl esterase activity (Biological processes, GO:0045330, *P*-value [adjusted] = 3.5×10^−2^), cell wall organization or biogenesis (GO:0071554, 4.548×10^−2^), and external encapsulating structure organization (GO:0045229, 6.873×10^−3^).

One of the genes affected by *ONSEN* was *FLC* (Chr5:3,173,042..3,180,975). This gene possesses a TEI in intron 1, which was previously identified as a hotspot for TEIs (Quadrana *et al*., 2016, 2019*b*; Baduel *et al*., 2021). Inspired by this observation, we analyzed all genes with *ONSEN* insertions in the current and a previously reported dataset (Baduel *et al*., 2021) and identified 29 such genes (Figure S3). Three of these genes were also found among the genes with *EVD* insertions according to (Baduel *et al*., 2021): the *FLC* locus, its antisense lncRNA *COOLAIR* (At5g01675) and the Class II formin gene *AtFH16* (At5g07770).

We tested the ability of CANS to detect transgenerationally inherited *ONSEN* insertions. We collected seeds (M1) from five M0 plants treated by heat stress and grown on medium containing α-amanitin and zebularine (Figure 5A). We grew 11 M1 plants (1 or 2 M1 plants per M0 plant) and performed CANS sequencing of a mixed DNA sample. In total, we detected 24 *ONSEN* insertions, including 16 supported by 2 or more reads. Some of the insertions were located in genes (Figure 5C) providing proof-of-concept that CANS is a useful tool for tracing individual TEIs after TE activation.

Altogether, using CANS, we were able to detect changes in the transposon activity of *ONSEN* family members in real time in response to heat stress. This information allowed us to identify biases in genome organization and hotspots of *ONSEN* insertions. CANS is also useful for detecting transgenerationally inherited TEIs, which could enable the rapid identification of the affected genes after TE activation in other plants including agronomically important crops.

### Detection of DNA methylation profiles of target TEs by CANS

DNA methylation on cytosine nucleotides is a pivotal modification for determining TE activity in plants (Fultz *et al*., 2015). The methylation patterns of ’young’ somatic TEIs are currently unclear and cannot be understood using short-read technologies (such as bisulfite sequencing). However, individual CANS reads contain cytosine methylation information that can be programmatically accessed (in this study, by DeepSignal-plant (Ni *et al*., 2021)). CANS reads that map to a single TE sequence could have originated from multiple somatic insertions. We reasoned that this information can be used to determine the average cytosine methylation profiles of somatic TEIs. We tested this assumption using the *ONSEN* transposon family and the CANS data collected from Col-0 plants (M0) grown on ZA medium and exposed to heat stress. We used CANS data obtained for M1 plants bearing new *ONSEN* insertions as a control. We successfully determined the high-resolution CG methylation profiles of the eight *ONSEN* family members (*ONSEN1–8*) using CANS data. We detected differences in CG methylation patterns among *ONSEN* family members and along TE sequences, with LTR sequences (400 bp) showing higher levels of methylation than the internal region (Figures S4-S9, Figure 6). Six *ONSEN* copies (*ONSEN2*, *ONSEN4-ONSEN8*) showed relatively similar CG methylation patterns between M0 and M1 plants according to this analysis (Figures S4–S9).

**Figure 6.**
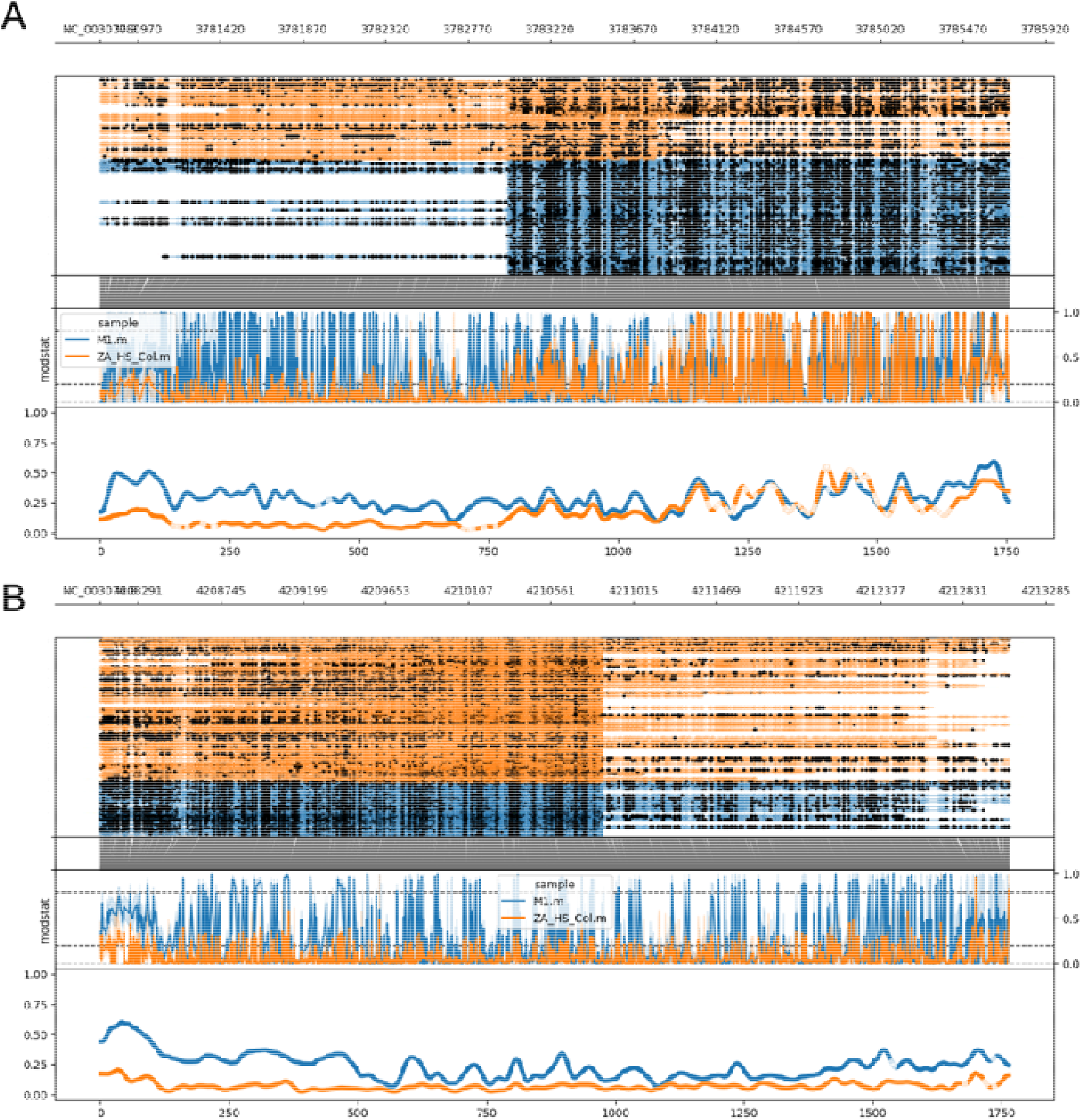
Distribution of CG methylation along ONSEN1 and ONSEN3 TEs deduced from CANS reads corresponding to reference and somatic TEIs. (A) *ONSEN1* (At1TE12295, NC_003070.9:3780765-3785721) and (B) *ONSEN3* (At5TE15240, NC_003076.8:4208083-4213084) read coverage plots and CG methylation patterns deduced by examining M0 (orange) and M1 (blue) CANS reads. From top to bottom: CANS read mapping showing methylated (closed circles) and unmethylated (opened circles) cytosine, raw log-likelihood ratios and smoothed methylated fraction plots.

Unexpectedly, *ONSEN1* and *ONSEN3* TEs showed significantly lower levels of CG methylation than the others (Figure 6). To determine the underlying cause, we assessed the transposition activity of *ONSEN* TEs. We used the number of reads with clipped ends corresponding to non-reference TE-genome junctions mapped to the individual *ONSEN*s as a proxy for transposition activity. According to this assay, transposition activity varied significantly among the eight *ONSEN*s, supporting an earlier report (Roquis *et al*., 2021). The most active *ONSEN* members in the Arabidopsis genome are *ONSEN1* (At1TE12295) and *ONSEN3* (At1TE59755), which together account for 61% of all *ONSEN* insertions.

Therefore, using CG methylation profiles inferred from CANS reads from individual TEs, we showed that *ONSEN1* and *ONSEN3* somatic insertions that were generated only a few days after heat stress tend to be hypomethylated. Overall, these results demonstrate the value of CANS data for exploring the cytosine methylation patterns of novel TEIs.

### *ONSEN* elements preferentially target genes downregulated by heat stress

The integration preferences of *ATCOPIA78* and *ATCOPIA93* have been linked to the histone variant H2A.Z (Quadrana *et al*., 2019; Roquis *et al*., 2021), which are found in genes that are sensitive to stress (Coleman-Derr and Zilberman, 2012). Based on previous data (Wollmann *et al*., 2017), we discovered that most (91%; 387) of the genes with somatic *ONSEN* insertions have higher than normal amounts of histone H2A.Z. However, the correlation between the expression levels of genes during heat stress and the rate of *ONSEN* insertions into these genes has not been explored. The clarify this point, we performed transcriptome deep sequencing (RNA-seq) analysis of Arabidopsis plants before (0-h samples) and 8, 16, or 24 h after heat stress (Figure 7A). We grew the plants in the ZA-containing medium described above in the same environment used to induce *ONSEN* insertions. Differential expression analysis of the 8-h, 16-h and 24-h samples vs. 0-h samples revealed 2327 (1214 upregulated and 1113 downregulated), 2675 (1379 upregulated and 1296 downregulated) and 2922 differentially expressed genes (DEGs; 1480 upregulated and 1442 downregulated), respectively (false discovery rate [FDR]< 0.01, fold-change > 2, *P*-value adjusted < 0.01) (Figure 7B, Supplemental Data S6-S8).

**Figure 7.**
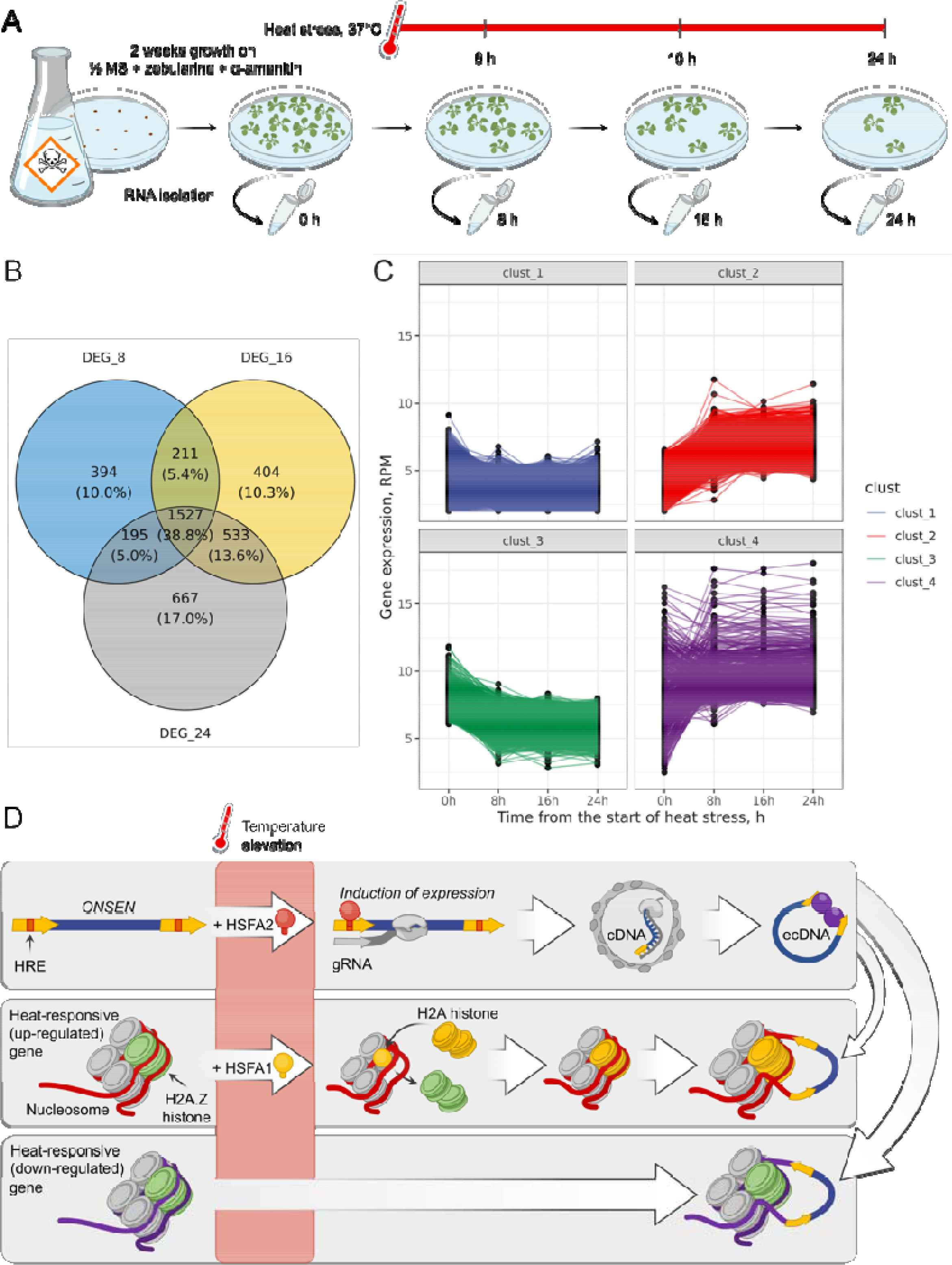
Transcriptome analysis of Arabidopsis plants grown on ZA-containing medium and subjected to heat stress. (A) Schematic overview of the RNA-seq experiment. (B) Venn diagram showing the number of upregulated and downregulated DEGs at 8, 16 and 24 h after the start of heat stress. (C) K-means clustering of DEGs. (D) A model of *ONSEN* insertions into genes that are downregulated in response to heat stress (37°C).

We compared these DEGs with the 426 genes containing *ONSEN* insertions (see above). The 8-h, 16-h, and 24-h DEGs included 47, 49, and 50 genes containing *ONSEN* insertions, respectively. The genes (32 of 426) with *ONSEN* insertions were significantly enriched in DEGs identified at all three time points (1527) under heat stress (Chi square contingency test *P*-value < 0.05). Surprisingly, gene set enrichment analysis revealed that genes that were downregulated at 8 h and 16 h of heat stress are enriched for *ONSEN* insertions (Chi square contingency test *P*-value 0.017 and 0.022, respectively), but we detected no enrichment for DEGs after 24 h of heat stress or for upregulated genes at any of the three time points. These findings indicate that downregulated genes are more likely than upregulated genes to acquire *ONSEN* insertions. This finding might be explained by the finding that histone H2A.Z is expelled from the nucleosome by HEAT SHOCK FACTOR A1 (HSFA1) and replaced by histone H2A (Cortijo *et al*., 2017). In the HSFA1 target gene set, we did not observe any enrichment or depletion of the genes with *ONSEN* insertions (Chi square contingency test *P*-value = 0.818).

To gain a deeper understanding of the relationship between *ONSEN* insertions and the dynamics of gene transcription under heat stress, we examined sets of genes with varying levels of dynamic transcription. Hierarchical clustering of 3931 DEGs revealed four major clusters of genes involved in the transcriptional response to heat stress (Figure 7C). Genes with minor changes in expression under heat stress form Cluster 1 (clust_1, 1547 genes). Genes with notably higher expression during heat stress define Cluster 2 (clust_2, 1035 genes). Genes that are highly expressed prior to heat stress (0 h) and significantly downregulated during heat stress are included in Cluster 3 (clust_3, 840 genes). The most heterogeneous cluster, Cluster 4 (clust_4, 509 genes), contains genes that exhibit various transcriptional patterns under heat stress. These four clusters contain 18, 19, 22, and 9 genes with detected ONSEN insertions, respectively. Enrichment analysis demonstrated that significantly more genes with *ONSEN* insertions are present in clust_3 than would be expected by chance (Chi square contingency test *P*-value = 0.001), supporting the link between the likelihood of an *ONSEN* insertion and the downregulation of gene expression under heat stress.

The combined data show that *ONSEN* has a propensity to insert into genes that are downregulated under heat stress. We propose that heat stress causes histone H2A.Z to be replaced by histone H2A at heat-responsive genes (upregulated during heat stress), reducing the likelihood that *ONSEN* will target these genes. Conversely, the downregulated genes continue to maintain histone H2A.Z, which attracts *ONSEN* insertions (Figure 7).

## Discussion

Here, we demonstrated that CANS is an easy-to-use, valuable tool for the rapid identification of T-DNA and TEIs in both repetitive (e.g. TE rich) and genic regions of plant genomes. CANS-based detection of T-DNA and TEI insertions has several advantages over current methods for TEI enrichment such as TE-capture (Quadrana *et al*., 2016). First, the CANS protocol, including ribonucleoprotein (RNP) complex formation, DNA cleavage and sequencing library preparation, is rapid and can be completed in 4-5 hours. Second, because a PCR amplification step is not needed, the resulting data can be used for epigenetic modification calling to generate methylation profiles of TEs or TEIs, although higher coverage is required for this purpose (Gabrieli *et al*., 2018; Ni *et al*., 2021). Third, long reads have high mappability to the genome, paving the way for detecting TEs and T-DNA insertions, even in repeat-rich regions of the genome such as centromeres and heterochromatin. For example, CANS revealed that many *EVD* insertions are located in pericentromeric regions and in other TEs. Fourth, NanoCasTE allows TEIs to be identified easily using CANS data, even if hundreds of insertions are captured. Finally, CANS is able to identify somatic mutations of target TEs, allowing rapid real-time monitoring of (i) the responses of the mobilome to stress, as we showed here using *ONSEN* and (ii) the insertion landscapes in different genetic backgrounds, as we demonstrated using *ddm1*. Despite the many advantages of CANS for TEI detection in plants, there are several issues that need to be overcome in the future: (1) a large amount of high-molecular-weight genomic DNA (>25 µg) is required; (2) deciphering the TE sequence of the insertion is challenging and requires the use of specific sets of overlapping sgRNAs; (3) only a small subset of sequencing pores (5-10%) perform sequencing at the same time, making the sequencing efficiency lower than that of WGS (90-98% of sequencing pores) and (4) the genotypes of TEIs (homozygous or hemizygous) cannot be determined using CANS.

While whole-genome Nanopore sequencing (WGNS) provides excellent resolution to potentially detect all TEIs in a genome (Kirov *et al*., 2021*a*; Roquis *et al*., 2021), it is expensive when examining a relatively large genome. For example, in our hands and based on previous reports using different species, it is challenging to generate more than 8 Gb of data per single MinION flow cell from plant material (Lee *et al*., 2019; Schalamun *et al*., 2019; Dmitriev *et al*., 2021). In contrast to WGNS, using only 4400 CANS reads (equivalent to 0.2× coverage of the Arabidopsis genome) on a single flow cell, we were able to detect all genetically inherited *EVD* insertions found by WGNS. We also unraveled the positions of T-DNA insertion sites using only one sgRNA. However, a higher number of sgRNAs and cleavage sites in genomic DNA may produce a higher proportion of target reads, as we observed in our pilot experiment with seven sgRNAs. The use of a higher number of sgRNAs per target is also thought to be beneficial for CANS of target genes (Gilpatrick *et al*., 2020). This is an important property of the CANS approach that allows more targets to be included per experiment without losing efficiency. Based on this assumption, insertions from many TEs would be expected to be captured in a single experiment using specifically designed and pooled sgRNAs. Here, we used multiplex CANS for the simultaneous detection of *EVD* and *ONSEN* insertions in Arabidopsis after heat stress. This approach was also recently used to detect TEIs in the human genome (McDonald *et al*., 2021).

We identified >800 TEIs in a small *ddm1* population after 2 hours of sequencing using only part of the capacity of MinION. This experiment provided the proof-of-concept that CANS is a valuable tool for identifying TEIs in a population. A study on the distribution of TEIs in the Arabidopsis 1001 Genomes panel demonstrated that TEs are a major source of large-effect alleles that can be fixed in populations, contributing to the local adaptation of different accessions (Baduel *et al*., 2021). Indeed, stress can induce the activation of TEs, which may lead to the rise of new alleles. TEI-derived alleles have spontaneously occurred in fields of crop species and have long been exploited in plant breeding programs (Kobayashi *et al*., 2004; Jiang *et al*., 2009; Butelli *et al*., 2012; Domínguez *et al*., 2020). Although the discovery of such alleles in crop fields using WGS is very challenging, CANS might represent a valuable alternative for capturing TEI-derived alleles in a population of cultivated plants. However, prior knowledge of active TEs that are capable of transposing is required for such experiments. Fortunately, this information is available for some plants. For example, we recently identified active LTR retrotransposons in sunflower (*Helianthus annuus*) (Kirov *et al*., 2020*b*) and triticale (× *Triticosecale Wittmack*) (Kirov *et al*., 2020*a*). There are many other examples of active plant TEs that can serve as potential targets for CANS-based capture of TEIs. The development of high-throughput tools that can capture TEIs in a population represents a milestone towards the acceleration of plant breeding and the more effective usage of natural variation. Further increasing the coverage of sequencing using tools such as the PromethION sequencer could greatly increase the sensitivity of this approach and might even enable the detection of TEIs that have occurred in a single plant within an entire population. Such results could then be used to design specific primers based on the insertion site in order to identify all plants carrying the target TEIs. The capture of TEIs occurring in individual plants in a population will also depend on the genome size of the species and the number of sgRNAs and target loci in the genome. These parameters must be optimized for individual plant species.

Estimating the integration preferences for distinct TEs is not straightforward, as post-integration selection pressure has a strong effect on the current TEI landscape. However, CANS is sensitive enough to capture somatic TEIs for which post-integration selection pressure has only a minor effect. This feature of CANS provides the opportunity to estimate the dynamic changes in the mobilome in real time. Therefore, CANS can uncover more TE target genes as insertions whose inheritance is deleterious can be detected. Here, we showed that *EVD* preferentially integrates into the pericentromeric regions of Arabidopsis chromosomes in the *ddm1* genome, whereas *ONSEN* transposons tended to integrate into genic regions and escape pericentromeric regions in the Arabidopsis Col-0 background. Notably, the established *EVD* insertion distribution was shaped by very recent events, with post-integration selection pressure having only a minor effect. Recently published data (Baduel *et al*., 2021) on the chromosomal distribution of TEIs in natural Arabidopsis accessions provide an excellent resource for comparing the genomic organizations of TEIs between natural and experimentally induced conditions.

The insertion preferences of *EVD*s in the *ddm1* mutant and natural Arabidopsis accessions (Baduel *et al*., 2021) and epiRILs (Quadrana *et al*., 2019) are not consistent: in the *ddm1* genome, *EVD* frequently targeted pericentromeric regions, while we identified almost no *EVD* insertions in these regions in the genomes of natural accessions and epiRILs. This discrepancy might be explained by differences in chromatin accessibility between wild-type and *ddm1* plants. Pericentromeric heterochromatin in more condensed in *ddm1* chromosomes than in the wild type (Probst *et al*., 2003), which can make these regions more attractive for *EVD* insertions in the mutant. Another possible explanation involves the differences in chromosomal distribution of histone H2A.Z. H2A.Z-containing nucleosomes are frequent targets of the *AtCOPIA93* retrotransposon family (Quadrana *et al*., 2019). However, this histone variant has an opposite distribution pattern from DNA methylation and is excluded from pericentromeric heterochromatin (Zilberman *et al*., 2008). Moreover, changes in DNA methylation (e.g. in *met1*) result in the redistribution of H2A.Z (Zilberman *et al*., 2008). Based on these observations, we suggest that the *ddm1* mutation causes H2A.Z to be loaded in hypomethylated pericentromeric regions, which stimulates the preferential insertion of *EVD* into these regions. Indeed, our analysis of *ONSEN* transposons showed that experimentally induced (via combined heat and epigenetic stress) and naturally occurring *ONSEN* insertions are well correlated at the genome level. Moreover, we identified 29 genes that were targets of *ONSEN* in both datasets, including *FLC*, pointing to the existence of hotspots for TEIs that result from TE integration preferences rather than from natural selection pressure.

The preferential targeting of certain chromatin states by TEs is thought to result in biased insertions into genes with specific expression patterns (Sultana *et al*., 2017). Such preferences were previously demonstrated for *Mutator* (*Mu*) transposons, which are frequently integrated into transcription start sites of genes expressed in the meristem (Zhang *et al*., 2020). The higher rate of TEIs into expressed loci is thought to be due to better accessibility of chromatin in the transcribed regions (Liu *et al*., 2009; Quadrana *et al*., 2019; Zhang *et al*., 2020). *ONSEN* transposons insert more frequently into genes associated with stress responses than others (Ito *et al*., 2016; Quadrana *et al*., 2019; Baduel *et al*., 2021b; Roquis *et al*., 2021). However, the connection between the expression patterns of genes during heat stress and the probability of *ONSEN* insertions has not yet been investigated. We showed that *ONSEN*s are more frequently inserted into genes that are downregulated at 8 and 16 h after the onset of heat stress. Heat stress has been shown to induce the expression of the heat shock factor protein, HSFA1, replacing H2A.Z nucleosomes at heat-responsive genes (upregulated under heat stress) with H2A histone (Cortijo *et al*., 2017). We propose that this replacement, in turn, makes these genes less attractive for *ONSEN* insertions compared to H2A.Z-associated genes (downregulated genes), making downregulated genes preferred targets for *ONSEN* insertions.

Our knowledge of the DNA methylation of TEs primarily comes from studies of genetically inherited insertions following meiosis. The fate of TEs immediately after insertion into the genome is poorly understood, as it requires analysis of somatic insertions, which is quite challenging. Oxford Nanopore reads and the associated raw signals provide a unique opportunity to drill deep into somatic insertion organization and DNA methylation. Here we used CANS reads to construct a high-density CG methylation profile of members of the *ONSEN* transposon family. *ONSEN* TEs are attractive for such studies, as their transposition can be triggered by heat stress and enhanced by DNA methylation and inhibited Polymerase II activity (Thieme *et al*., 2017). CANS sequencing of *ONSEN*s following the induction of their transposition allows reads to be obtained from the donor (reference) TE loci as well as from somatic insertions. We exploited this system to explore the methylation patterns of *ONSEN* somatic insertions after a few days of heat stress. We found that the distribution of CG methylation differs among *ONSEN* family members and that the level of DNA methylation at somatic insertions is significantly reduced for two of the most active *ONSEN* members (*ONSEN1* and *ONSEN3*). The hyperactivity of the ONSEN elements may interfere with the establishment of DNA methylation at new insertions. Alternatively, the positions of novel insertions may differ among *ONSEN* family members, which influences DNA methylation. CANS sequencing with a significantly higher sequencing depth for somatic insertions (e.g. using PromethION) is needed to examine this issue.

Overall, our work demonstrates that CANS is a valuable and cost-effective approach to rapidly decipher the mobilome landscapes of plant genomes and to elucidate changes in DNA methylation in TEs in plants with different genetic backgrounds and under stress conditions.

## Material and Methods

### Plant material and growth conditions

Seeds of ddm1 mutants (ddm1-2, F7 generation) were kindly provided by Vincent Colot (Institut de Biologie de l’Ecole Normale Supérieure (IBENS), Paris, France). Arabidopsis thaliana Col-0 plants (wild type and ddm1 mutants) were grown in a light chamber for a month under 22°C and long-day conditions (16h light/8h dark). For sequencing of pooled plants, Arabidopsis seeds were surface-sterilized with 75% ethanol for 2 min, then washed with 5% sodium hypochlorite for 5 min. After this, the seeds were rinsed with sterile distilled water 3 times. 0,1% agarose was added to each tube and the seeds were resuspended and dripped on Petri dishes with MS nutrient medium (Murashige and Skoog, 1962), supplemented with 3% of sucrose (PanReac AppliChem, Germany) and 1% of agar (PanReac AppliChem, Germany). Plates with seeds were sealed with Parafilm (Pechiney Plastic Packaging Company, USA) and kept in the dark at 4°C for 3 days for vernalization and synchronous germination. Afterwards, dishes were transferred into a light chamber with 16h day/8h night photoperiods for further growth.

### ONSEN activation in vitro

The principle of heat mediated activation of *ONSEN* has been previously described (Thieme *et al*., 2017). For aseptic growth, seeds of *A. thaliana* were sterilized in 10% bleach for 10 min, rinsed three times with sterile H2O, and sown on 9 cm Petri dishes containing half-strength MS media (1% sucrose, 0.6% agar (Dia-M, 3440.0500), pH 5.7) supplemented with combination of sterile filtered 5 µM α-amanitin (dissolved in H2O) and 40 µM zebularine (dissolved in DMSO) (Thieme *et al*., 2017). After sowing the plates were placed at 4℃ in the dark for 2 days. Germinated seedlings were then grown in a growth chamber (23℃) under long-day conditions (16:8h, light:dark). After 10 days, the plantlets were put at 6◦C for 24 h (lighting condition unchanged). The plants were then submitted to 24 h of heat stress at 37℃ (HS). Then transferred to long day conditions (23°C, 16h/8h light/dark photoperiod).

For seed collection from the plants after *ONSEN* activation the plants were grown according to the previously described protocol (Pasternak *et al*., 2020). Briefly, plants were transferred from α-amanitin/zebularine containing medium to TK1 medium in 1% agar and kept in containers with TK1 until plants transit to bolting. Further, shoots were grown on Hg medium until seed formation.

### Plant transformation

To create ddm1 plant carrying T-DNA od nucleotide sequence of the *GAG* open reading frame of *EVD5* retrotransposon (*AtCOPIA93* (Mirouze *et al*., 2009)). The *GAG* ORF sequence was synthesized by the Synbio Technologies service (USA). The *GAG* fragment was cloned at the restriction sites BamH1 and Sac1 into the pBI121 vector. The Agrobacterium transformation of *A. thaliana* plants was carried out as described previously (Clough and Bent, 1998). Primary transformants were selected based on their resistance to kanamycin (50 lg/ml). At least 60 transgenic seeds of the T2 generation were selected and used for segregation analysis. A plant from one T2 family (G-ddm1-3) with 3:1 segregation of the marker for kanamycin resistance was then transferred to pots with soil, grown three weeks more and used for DNA isolation for whole-genome and CANS sequencing.

### HMW DNA isolation and size selection

High molecular weight DNA was isolated from 200-500 mg of fresh and young leaves that were homogenized in liquid nitrogen. DNA isolation was carried out according to the previously published protocol (https://www.protocols.io/view/plant-dna-extraction-and-preparation-for-ont-seque-bcvyiw7w) with several modifications. Chloroform was used instead of dichloromethane during the extraction step. Samples with added CTAB2 buffer were incubated at 65°С until the formation of visible flakes. Dissolving of the precipitate in 1M NaCl was performed at 65°С for 15 minutes with further cooling to the RT before adding isopropanol. Removal of contaminants residues and small-size nucleotide fragments was performed by adding 2.5 volumes of 80% ethanol with further centrifugation at 12000 g for 15 minutes. The obtained DNA pellet was washed with 70% ethanol and centrifuged at 12000 g for 5 minutes. Final DNA elution was carried out with 65 µL of nuclease-free water.

For whole-genome Nanopore sequencing and CANS, size-selection of the large DNA fragments was done using SRE or XL Short Read Eliminator Kits (Circulomics, USA) according to the manufacture instructions.

The concentration and quality of isolated DNA were estimated by gel electrophoresis in 1% agarose gel, NanoDrop One UV-Vis Spectrophotometer (Thermo Scientific, USA) and Quantus Fluorometer (Promega, USA) using a DNA QuantiFluor ONE dsDNA System (Promega, USA). Only pure DNA (A260/A280 ∼1.8 and A260/A230 ∼2.0 according to NanoDrop) with almost no differences in concentrations obtained by Nanodrop and Quantus was used for sequencing.

### sgRNA design

To design sgRNAs recognizing multiple copies of LTR retrotransposons we used a step-by-step strategy: (1) using FlashFry ‘discover’ tool with the ‘-positionOutput -maxMismatch 0 ‘ arguments and reference genome sequences (TAIR10) a pool of sgRNAs and the corresponding genomic sites were found followed by bed file generation; (2) reference sequences of target LTR retrotransposons were blasted against the corresponding genomes followed by hit filtering (minimum hit length 100bp and minimum e-value 1e-50) and bed file consisting blast hit positions was generated; (3) the genomic location of predicted sgRNA sites (step 1) were intersected with blast hit bed file (step 2) using bedtools ’intersect’ and OnOutTE ratio was calculated for each sgRNAs.

OnOutTE = number of sgRNA sites located on the blast hit from the corresponding TE / total number of sgRNA sites

(4) We selected sgRNAs with OnOutTE ratio > 0.85.

For each retrotransposon a set of sgRNAs were designed including one (*A. thaliana*) sgRNAs on negative strand recognizing both LTRs and three sgRNAs (one on negative and two on positive strands) located between two LTR sequences. Sequences of sgRNAs are provided in Supplemental Data S1.

### In vitro transcription of sgRNAs

SgRNAs for CANS were produced by in vitro transcription from double-stranded DNA templates containing T7 promoter that were assembled from two oligos, specific (sgRNA1-5 in Supplemental Data S1) and universal (CRISPR_R). All oligonucleotides were ordered in Evrogen (Moscow, Russia). To synthesize double-stranded DNA templates, the following reagents were mixed per each reaction: 2μl of unique sgRNA oligo (1 μM); 2μl of CRISPR_R (1 μM); 2μl of T7 forward (5’-GGATCCTAATACGACTCACTATAG-3’) and reverse (5’-AAAAAAGCACCGACTCGG-3’) primers (100 μM of each); 2μl of 50x dNTP (10 μM) (Evrogen, Moscow, Russia); 10μl of 10x Encyclo buffer (Evrogen, Moscow, Russia); 1μl of Encyclo polymerase (Evrogen, Moscow, Russia); 79μl of RNAse-free water. Oligos were annealed and extended according to the PCR program: initial denaturation at 95°C (2 min); 30 cycles of denaturation at 98°C (30 sec), annealing at 60°C (30 sec), elongation at 72°C (30 sec); final elongation at 72°C (1 min). Gel electrophoresis was performed after amplification and templates were column purified.

To transcribe the templates, the Highly Efficient RNA in vitro Synthesis Kit (Biolabmix, Novosibirsk, Russia) was used. The subsequent reactions were set up and incubated at 37°C for 2h: 10μl of 5x T7 transcription buffer; 2μl of 25x DTT (250mM); 2μl of dNTP (25mM of each); 2μl of double-stranded DNA template (150-200 ng); 1μl of T7 polymerase (150 units); 33μl RNAse-free water. The sgRNAs were then purified using RNA and miRNA Extraction Kit (Biolabmix, Novosibirsk, Russia) according to the manufacturer instructions. The concentration and quality of prepared sgRNAs were estimated by Nanodrop (Thermo Scientific, USA), Qubit (Thermo Scientific, USA) and gel electrophoresis in 2% agarose gel.

### Cas9/sgRNA ribonucleoprotein complexes (RNPs) assembly

For CANS, 50 ng of each sgRNA were used for RNP assembly. To obtain RNPs for sgRNA Mixture 2 the corresponding sgRNAs were pooled together at 1:1 molar ratio in 11 μl of MQ water. Before RNP assembly all sgRNA mixtures were denaturated at 95°C for 5 min, then cooled on ice. The RNPs were assembled by combining 11μl of sgRNA mix with 1μl of Cas9 nuclease in 3 μl of reaction buffer (Biolabmix, Novosibirsk, Russia) at a final volume of 15μl followed by incubation for 30 min at room temperature and kept on ice until usage.

### Cas9 cleavage and library preparation

Before CANS DNA was dephosphorylated. For this, 2-8 µg of HMW DNA was diluted in 1x CutSmart Buffer and 6 µl of Quick CIP enzyme (New England Biolabs, catalog no. M0508) was added followed by incubation of the reaction at 37°C for 30 minutes. The reaction was stopped by heating (80°C, 2 minutes). DNA cleavage by RNPs was carried out by mixing of 40 µl of dephosphorylated DNA, 15 µl of RNPs, 1.5 µl of dATP (10 mM) and 1 µl Encyclo polymerase (Evrogen, Moscow, Russia). The mixture was incubated at 37°C for 30 minutes followed by 5□min at 72□°C. Nanopore sequencing adapters (AMX) were ligated to the Cas9 cleaved DNA by mixing the following components: 25 µl of LNB buffer (Oxford Nanopore Technologies, catalog no. SQK-LSK109), 5 µl of nuclease-free water, 12.5 µl of Quick T4 DNA Ligase (New England Biolabs, NEBNext Companion Module for Oxford Nanopore Technologies Ligation Sequencing catalog no. E7180S) and Adapter Mix (SQK-LSK109). The mixture was added to the Cas9 cleaved DNA and incubated for 20 min at room temperature. The samples were purified by adding equal volume of TE buffer (pH 8.0) and 0.3 volume of AMPure XP Beads (Beckman Coulter, catalog no. A63881) and washed twice by LFB buffer (Oxford Nanopore Technologies, catalog no. SQK-LSK109). The DNA was eluted in 15□μl of elution buffer (Oxford Nanopore Technologies, catalog no. SQK-LSK109). Then sequencing library was prepared according to the Ligation Sequencing Kit SQK-LSK109 protocol (Oxford Nanopore Technologies, catalog no. SQK-LSK109).

Library preparation for whole-genome sequencing was carried out using the Ligation Sequencing Kit (Oxford Nanopore Technologies, catalog no. SQK-LSK109) with 1 µg of input HMW DNA.

### Sequencing and basecalling

Sequencing was performed by MinION equipped by R9.4.1 or R10.3 flow cells. Sequencing process was operated by MinKNOW software (v.19.12.5). The detailed information on Nanopore sequencing is described in Supplemental Data S2.

### Analysis

Base calling was done by Guppy (Version 3.2.10). The TAIR10 genome was downloaded from NCBI database. The generated Nanopore reads were aligned to the reference genomes (TAIR10 for *A.thaliana*) using minimap2 software (Li, 2018) with the following parameters: -ax map-ont -t 100. The obtained sam file was converted to bam format, sorted and indexed using SAMtools (Li *et al*., 2009). The obtained sorted bam files were used for NanoCasTE (CANS data) and nanotei (WGS data) pipelines.

### NanoCasTE pipeline for the identification of transposon insertions following CANS

NanoCasTE is written in python and can be run as a standalone tool or via Snakefile (https://github.com/Kirovez/NanoCasTE). NanoCasTE involves several steps (Figure 8): (1) Mapping the reads to the genome using minimap2 (Li, 2018), followed by the generation of sorted bam files using SAMtools (Li *et al*., 2009); (2) Filtering the reads based on mapping quality (MQ > 40), read length (>750) and SA tags (no supplementary alignments are allowed); (3) Selecting mapped reads with clipped heads (reads mapped to the positive strand) or tails (reads mapped to the negative strand) with the length of the clipped parts close to the distance from the sgRNA positions to the TE end; (4) Filtering the reads based on similarity of the clipped part to the target TE sequence; (5) Identifying the inserted sites and TE orientation (+ or - strand) and outputting the results.

**Figure 8.**
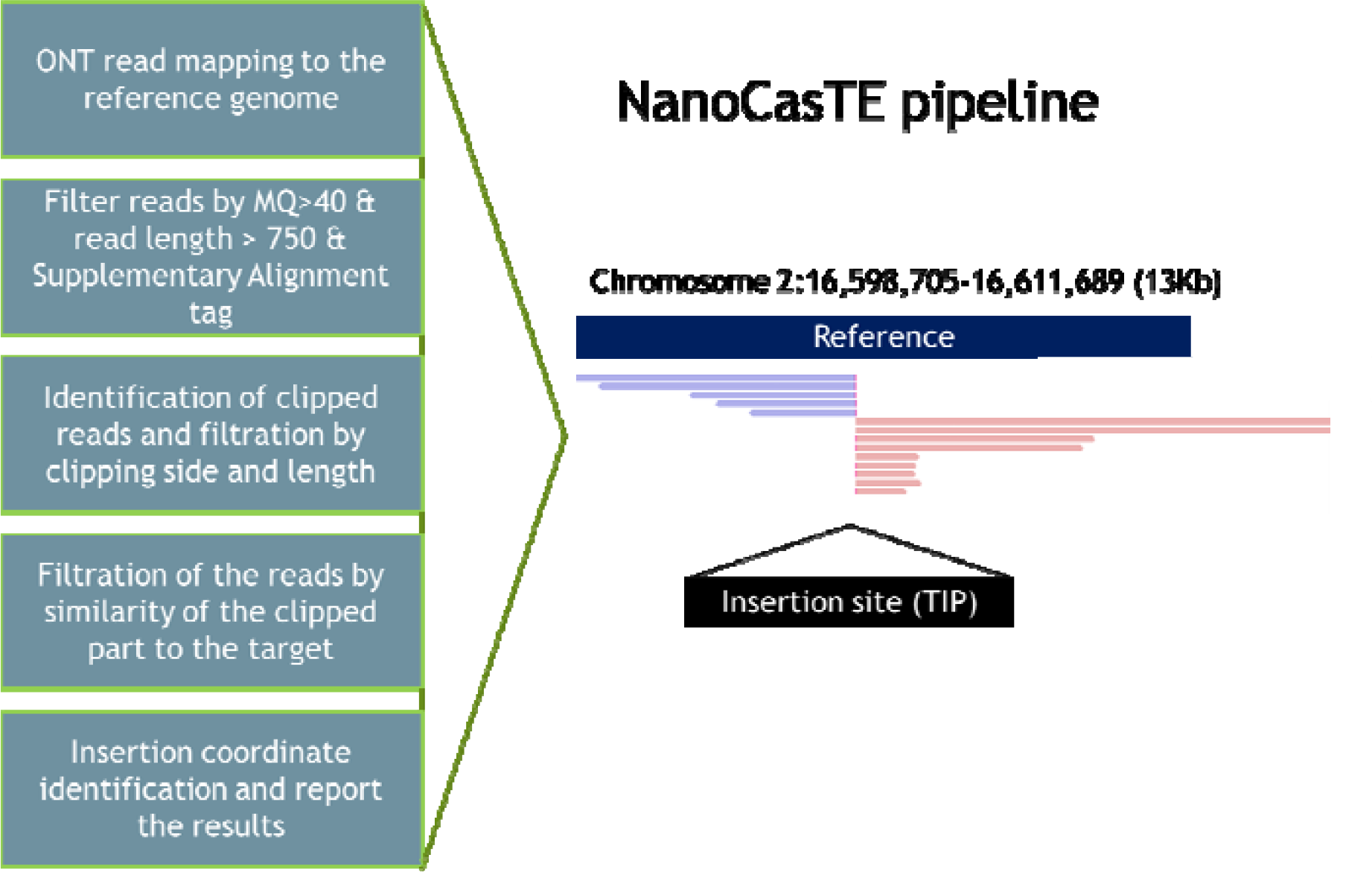
Schematic view of the major steps of the NanoCasTE pipeline to identify transposon insertions in the genome after CANS.

NanoCasTE uses a set of stringent criteria to specifically detect TE insertions in CANS data and to distinguish them from noise signals. The pipeline reports the putative positions of the target TE insertion as well as additional information that is useful for further analysis, including the number of selected reads supporting the insertion, the total number of reads covering the insertion, the strand harboring the TE insertion and the length of the clipped parts of the selected reads.

NanoCasTE was run for all samples using the following parameters: min_len_clipped: 0.3, mapping_quality: 60, min_read_length: 700. The following lists of sgRNAs were used for *A. thaliana EVD:* ’TCTTGGTGATGAGAGTGAC, ACCCTGGATTTAAGGTGAGA, AGTTTAAGAGCTCTAGTATG, CTACAAGGTCAATCGAAAGG, TCAACACATGAAAGTCCCGA’; for *A. thaliana* ONSEN: ′GTTTAAAGCCCATGTTGAGA,TCAAGGAAGGAAATGGTGAG,ACCGTTTCTAGACGAGCAAC,TCGAGTATTCCAAGCTCTTG′.

## Methylation Calling

Methylation calling from raw Nanopore data was performed using DeepSignal-plant software (Ni *et al*., 2021) as previously described (Kirov *et al*., 2021*b*). For methylation visualization, the custom-made script DeepS2bam_converter (https://github.com/Kirovez/DeepS2bam_converter, accessed on 10 March 2021) was utilized to manually check the CG methylation distribution. The read alignments and per-read methylation profile were visualized by JBrowse2 (Buels *et al*., 2016). Methylartist pipeline was used for CG methylation plot drawing (Cheetham *et al*., 2022).

### Insertion validation by PCR

For validations of TEIs we used the combination of primers including one primer located on TEI flanking regions and one primer located on TE. The sequences of primers are listed in Supplemental Data S3. The PCR was performed in a reaction volume of 25 µl using Bio-Rad Т100 Thermal Cycler (Bio-Rad, USA). The reaction mixture contained 2.5 mM MgCl2, 200 mM dNTPs (Dia-M, Moscow, Russia), 5 pmols of each primer, 0.5 units of ColoredTaq polymerase (Sileks, Russia). The PCR reactions were carried out under the following conditions: Touchdown-PCR was carried out according to the following cycling program: 94 ◦C for 3 min, 94 ◦C for 30 s, 65 ◦C for 30 s and 72 ◦C for 2 min, followed by 6 cycles at decreasing annealing temperatures in decrements of 1 ◦C per cycle, then 35 cycles of 30 s at 94 ◦C, 30 s at 55 ◦C, 2 min at 72 ◦C and final extension at 72◦C for 5 min. For validation of *A. thaliana* TEIs the following PCR program was used: 35 cycles of 94◦C for 30 s, 60◦C for 30 s and 72 ◦C for 2 min. Amplification products were visualized by electrophoresis in 1.5% agarose gel and observed under UV light after ethidium bromide staining.

### RNAseq analysis of transcriptome changes during heat stress

For RNA sequencing arabidopsis Col-0 plants were grown on 1/2 MS medium supplemented with combination of sterile filtered 5 µM α-amanitin (dissolved in H2O) and 40 µM zebularine (dissolved in DMSO) (Thieme *et al*., 2017). After sowing the plates were placed at 4℃ in the dark for 2 days. Germinated seedlings were then grown in a growth chamber (23℃) under long-day conditions (16:8h, light:dark). After 10 days, the plantlets were put at 6◦C for 24 h (lighting condition unchanged). The plants were then submitted to 24 h of heat stress at 37℃ (HS). RNA was isolated from two biological replicates each for 4 time points corresponding to 0 (control) and 8, 16 and 24 hours after heat stress starts. RNA was isolated using the ExtractRNA kit (Evrogen, Moscow, Russia) according to the manufacturer’s instructions. mRNA sequencing was performed with assistance of BGI (Shenzhen, China) on DNBSEQ (DNBSEQ Technology) platform. The raw reads were pre-processed using BGI-Tech bioinformatics workflow. For each sample >36M clean paired-end 100bp reads have been obtained and utilized for deferential expression analysis. Reads were were mapped to TAIR10 genome using HISAT2 (Kim *et al*., 2019) with “-k 2” parameter. Htseq-count (Anders *et al*., 2015) was used to assign reads to exons with the following settings: “--stranded yes --minaqual 20 --type exon --idattr gene_id -m union”. Quality control, normalization, and scaling of the raw read counts were performed by EdgeR (Robinson *et al*., 2010) package as a part of iDEP server (Differential Expression and Pathway analysis (iDEP.95)) (Ge *et al*., 2018). Differentially expressed genes were identified using the following criteria: fold-change > 2, P-value (adjusted) < 0.01, FDR cutoff = 0.01.

### Statistics and Data Visualization

Statistical analysis was carried out in Rstudio Version 1.2.1335 (http://www.rstudio.com/) with R version 3.6.0. Visualization was carried out by ggplot2 (Wickham, 2011) and VennDiagram (Chen and Boutros, 2011) packages. The circus plots were drawn using Circos software (Krzywinski *et al*., 2009) implemented in Galaxy server (Rasche and Hiltemann, 2020).

### Accession Numbers

All Nanopore and RNA-seq data generated in this study were deposited in the National Center for Biotechnology Information SRA database (BioProject ID PRJNA736208).

## Acknowledgements

We thank Vincent Colot (Institut de Biologie de l’Ecole Normale Supérieure (IBENS), Paris, France) for providing the *ddm1* plant materials.

## Author contribution statement

I.K., P.M. and M.D. designed the research; I.K., P.M., S.G., R.K., M.O., M.D., Z.K. and A.K. performed the research. I.K., G.K., P.M., A.S. and M.D. analyzed the data. I.K. and P.M. wrote the article.

## References

Anders S, Pyl PT, Huber W. 2015. HTSeq—a Python framework to work with high-throughput sequencing data. Bioinformatics 31, 166–169.

Baduel P, Leduque B, Ignace A, Gy I, Gil J, Loudet O, Colot V, Quadrana L. 2021. Genetic and environmental modulation of transposition shapes the evolutionary potential of Arabidopsis thaliana. Genome Biology 22, 138.

Buels R, Yao E, Diesh CM, et al. 2016. JBrowse: a dynamic web platform for genome visualization and analysis. Genome Biology 17, 66.

Butelli E, Licciardello C, Zhang Y, Liu J, Mackay S, Bailey P, Reforgiato-Recupero G, Martin C. 2012. Retrotransposons control fruit-specific, cold-dependent accumulation of anthocyanins in blood oranges. Plant Cell 24, 1242–55.

Carpentier MC, Manfroi E, Wei FJ, Wu HP, Lasserre E, Llauro C, Debladis E, Akakpo R, Hsing YI, Panaud O. 2019. Retrotranspositional landscape of Asian rice revealed by 3000 genomes. Nat Commun 10, 24.

Cheetham SW, Kindlova M, Ewing AD. 2022. Methylartist: Tools for Visualising Modified Bases from Nanopore Sequence Data. Bioinformatics 38, btac292-.

Chen H, Boutros PC. 2011. VennDiagram: a package for the generation of highly-customizable Venn and Euler diagrams in R. BMC Bioinformatics 12, 35.

Chen J, Lu L, Robb SMC, Collin M, Okumoto Y, Stajich JE, Wessler SR. 2020. Genomic diversity generated by a transposable element burst in a rice recombinant inbred population. Proc Natl Acad Sci U S A 117, 26288–26297.

Chuong EB, Elde NC, Feschotte C. 2017. Regulatory activities of transposable elements: from conflicts to benefits. Nature Reviews Genetics 18, 71–86.

Clough SJ, Bent AF. 1998. Floral dip: a simplified method for Agrobacterium-mediated transformation of Arabidopsis thaliana. Plant J 16, 735–43.

Coleman-Derr D, Zilberman D. 2012. Deposition of Histone Variant H2A.Z within Gene Bodies Regulates Responsive Genes. PLoS Genetics 8, e1002988.

Cortijo S, Charoensawan V, Brestovitsky A, Buning R, Ravarani C, Rhodes D, Noort J van, Jaeger KE, Wigge PA. 2017. Transcriptional Regulation of the Ambient Temperature Response by H2A.Z Nucleosomes and HSF1 Transcription Factors in Arabidopsis. Molecular Plant 10, 1258–1273.

Dmitriev AA, Pushkova EN, Novakovskiy RO, et al. 2021. Genome Sequencing of Fiber Flax Cultivar Atlant Using Oxford Nanopore and Illumina Platforms. Frontiers in genetics 11, 590282–590282.

Domínguez M, Dugas E, Benchouaia M, Leduque B, Jiménez-Gómez JM, Colot V, Quadrana L. 2020. The impact of transposable elements on tomato diversity. Nature Communications 11, 4058.

Fultz D, Choudury SG, Slotkin RK. 2015. Silencing of active transposable elements in plants. Current Opinion in Plant Biology 27, 67–76.

Gabrieli T, Sharim H, Fridman D, Arbib N, Michaeli Y, Ebenstein Y. 2018. Selective nanopore sequencing of human BRCA1 by Cas9-assisted targeting of chromosome segments (CATCH). Nucleic Acids Research 46, e87–e87.

Ge SX, Son EW, Yao R. 2018. iDEP: an integrated web application for differential expression and pathway analysis of RNA-Seq data. BMC Bioinformatics 19, 534.

Gilpatrick T, Lee I, Graham JE, Raimondeau E, Bowen R, Heron A, Downs B, Sukumar S, Sedlazeck FJ, Timp W. 2020. Targeted nanopore sequencing with Cas9-guided adapter ligation. Nature Biotechnology 38, 433–438.

Handsaker RE, Korn JM, Nemesh J, McCarroll SA. 2011. Discovery and genotyping of genome structural polymorphism by sequencing on a population scale. Nat Genet 43, 269–76.

Ito H, Gaubert H, Bucher E, Mirouze M, Vaillant I, Paszkowski J. 2011. An siRNA pathway prevents transgenerational retrotransposition in plants subjected to stress. Nature 472, 115–119.

Ito H, Kim J-M, Matsunaga W, et al. 2016. A Stress-Activated Transposon in Arabidopsis Induces Transgenerational Abscisic Acid Insensitivity. Scientific Reports 6, 23181.

Jiang N, Gao D, Xiao H, Knaap E van der. 2009. Genome organization of the tomato sun locus and characterization of the unusual retrotransposon Rider. Plant J 60, 181–93.

Jung H, Winefield C, Bombarely A, Prentis P, Waterhouse P. 2019. Tools and Strategies for Long-Read Sequencing and De Novo Assembly of Plant Genomes. Trends in Plant Science 24, 700–724.

Kim D, Paggi JM, Park C, Bennett C, Salzberg SL. 2019. Graph-based genome alignment and genotyping with HISAT2 and HISAT-genotype. Nature Biotechnology 37, 907–915.

Kirov I, Dudnikov M, Merkulov P, Shingaliev A, Omarov M, Kolganova E, Sigaeva A, Karlov G, Soloviev A. 2020a. Nanopore RNA Sequencing Revealed Long Non-Coding and LTR Retrotransposon-Related RNAs Expressed at Early Stages of Triticale SEED Development. Plants 9, 1794.

Kirov I, Merkulov P, Dudnikov M, et al. 2021a. Transposons Hidden in Arabidopsis thaliana Genome Assembly Gaps and Mobilization of Non-Autonomous LTR Retrotransposons Unravelled by Nanotei Pipeline. Plants 10, 2681.

Kirov I, Omarov M, Merkulov P, Dudnikov M, Gvaramiya S, Kolganova E, Komakhin R, Karlov G, Soloviev A. 2020b. Genomic and Transcriptomic Survey Provides New Insight into the Organization and Transposition Activity of Highly Expressed LTR Retrotransposons of Sunflower (Helianthus annuus L.). Int J Mol Sci 21, 9331.

Kirov I, Polkhovskaya E, Dudnikov M, Merkulov P, Vlasova A, Karlov G, Soloviev A. 2021b. Searching for a Needle in a Haystack: Cas9-Targeted Nanopore Sequencing and DNA Methylation Profiling of Full-Length Glutenin Genes in a Big Cereal Genome. Plants 11, 5.

Kobayashi S, Goto-Yamamoto N, Hirochika H. 2004. Retrotransposon-induced mutations in grape skin color. Science 304, 982.

Krzywinski M, Schein J, Birol İ, Connors J, Gascoyne R, Horsman D, Jones SJ, Marra MA. 2009. Circos: An information aesthetic for comparative genomics. Genome Research 19, 1639–1645.

Lee YG, Choi SC, Kang Y, Kim KM, Kang C-S, Kim C. 2019. Constructing a Reference Genome in a Single Lab: The Possibility to Use Oxford Nanopore Technology. Plants 8, 270.

Li H. 2018. Minimap2: pairwise alignment for nucleotide sequences. Bioinformatics 34, 3094–3100.

Li H, Handsaker B, Wysoker A, Fennell T, Ruan J, Homer N, Marth G, Abecasis G, Durbin R, Processing SGPD. 2009. The Sequence Alignment/Map format and SAMtools. Bioinformatics 25, 2078–9.

Li S, Jia S, Hou L, Nguyen H, Sato S, Holding D, Cahoon E, Zhang C, Clemente T, Yu B. 2019. Mapping of transgenic alleles in soybean using a nanopore-based sequencing strategy. Journal of Experimental Botany 70, 3825–3833.

Lisch D. NRG.. 2013. How important are transposons for plant evolution? Nature Reviews Genetics 14, 49–61.

Liu S, Yeh C-T, Ji T, Ying K, Wu H, Tang HM, Fu Y, Nettleton D, Schnable PS. 2009. Mu Transposon Insertion Sites and Meiotic Recombination Events Co-Localize with Epigenetic Marks for Open Chromatin across the Maize Genome. PLoS Genetics 5, e1000733.

López-Girona E, Davy MW, Albert NW, Hilario E, Smart MEM, Kirk C, Thomson SJ, Chagné D. 2020. CRISPR-Cas9 enrichment and long read sequencing for fine mapping in plants. Plant Methods 16, 121.

Madsen EB, Höijer I, Kvist T, Ameur A, Mikkelsen MJ. 2020. Xdrop: Targeted sequencing of long DNA molecules from low input samples using droplet sorting. Hum Mutat 41, 1671–1679.

McDonald TL, Zhou W, Castro C, Mumm C, Switzenberg JA, Mills RE, Boyle AP. 2021. Cas9 targeted enrichment of mobile elements using nanopore sequencing., 2021.02.10.430605.

Mirouze M, Reinders J, Bucher E, Nishimura T, Schneeberger K, Ossowski S, Cao J, Weigel D, Paszkowski J, Mathieu O. 2009. Selective epigenetic control of retrotransposition in Arabidopsis. Nature 461, 427–30.

Ni P, Huang N, Nie F, et al. 2021a. Genome-wide Detection of Cytosine Methylations in plant from Nanopore data using Deep Learning. Nature Communications 12, 5976.

et al.et al.Nuthikattu S, McCue AD, Panda K, Fultz D, DeFraia C, Thomas EN, Slotkin RK. 2013. The Initiation of Epigenetic Silencing of Active Transposable Elements Is Triggered by RDR6 and 21-22 Nucleotide Small Interfering RNAs. 162, 116–131.

Panda K, Slotkin RK. 2020. Long-Read cDNA Sequencing Enables a “Gene-Like” Transcript Annotation of Transposable Elements. The Plant Cell 32, 2687–2698.

Pasternak T, Ruperti B, Palme K. 2020. A simple high efficiency and low cost in vitro growth system for phenotypic characterization and seed propagation of Arabidopsis thaliana. bioRxiv, 2020.08.23.263491.

Probst AV, Fransz PF, Paszkowski J, Scheid OM. 2003. Two means of transcriptional reactivation within heterochromatin. The Plant Journal 33, 743–749.

Quadrana L, Etcheverry M, Gilly A, et al. 2019. Transposition favors the generation of large effect mutations that may facilitate rapid adaption. Nature Communications 10, 3421.

et al.Quadrana L, Silveira AB, Mayhew GF, LeBlanc C, Martienssen RA, Jeddeloh JA, Colot V. 2016b. The Arabidopsis thaliana mobilome and its impact at the species level. eLife 5, e15716.

Rabanus-Wallace MT, Hackauf B, Mascher M, et al. 2021. Chromosome-scale genome assembly provides insights into rye biology, evolution and agronomic potential. Nature Genetics 53, 564–573.

Rasche H, Hiltemann S. 2020. Galactic Circos: User-friendly Circos plots within the Galaxy platform. GigaScience 9, giaa065-.

Robinson MD, McCarthy DJ, Smyth GK. 2010. edgeR: a Bioconductor package for differential expression analysis of digital gene expression data. Bioinformatics 26, 139–140.

Roquis D, Robertson M, Yu L, Thieme M, Julkowska M, Bucher E. 2021. Genomic impact of stress-induced transposable element mobility in Arabidopsis. Nucleic Acids Research 49, 10431–10447.

Schalamun M, Nagar R, Kainer D, Beavan E, Eccles D, Rathjen JP, Lanfear R, Schwessinger B. 2019. Harnessing the MinION: An example of how to establish long-read sequencing in a laboratory using challenging plant tissue from Eucalyptus pauciflora. Mol Ecol Resour 19, 77–89.

Slotkin RK, Martienssen R. 2007. Transposable elements and the epigenetic regulation of the genome. Nature Reviews Genetics 8, 272–285.

Stangl C, Blank S de, Renkens I, et al. 2020. Partner independent fusion gene detection by multiplexed CRISPR-Cas9 enrichment and long read nanopore sequencing. Nature Communications 11, 2861.

Sultana T, Zamborlini A, Cristofari G, Lesage P.J NRG. 2017a. Integration site selection by retroviruses and transposable elements in eukaryotes. Nature Reviews Genetics 18, 292–308.

Thieme M, Lanciano S, Balzergue S, Daccord N, Mirouze M, Bucher E. 2017. Inhibition of RNA polymerase II allows controlled mobilisation of retrotransposons for plant breeding. Genome Biology 18, 134.

Tsukahara S, Kobayashi A, Kawabe A, Mathieu O, Miura A, Kakutani T. 2009. Bursts of retrotransposition reproduced in Arabidopsis. Nature 461, 423–6.

Wickham H. 2011. ggplot2. Wiley Interdisciplinary Reviews: Computational Statistics 3.2 3, 180–185.

Williams-Carrier R, Stiffler N, Belcher S, Kroeger T, Stern DB, Monde R-A, Coalter R, Barkan A. 2010. Use of Illumina sequencing to identify transposon insertions underlying mutant phenotypes in high-copy Mutator lines of maize. Plant J 63, 167–177.

Wollmann H, Stroud H, Yelagandula R, et al. 2017. The histone H3 variant H3.3 regulates gene body DNA methylation in Arabidopsis thaliana. Genome Biology 18, 94.

Zhang X, Zhao M, McCarty DR, Lisch D. 2020. Transposable elements employ distinct integration strategies with respect to transcriptional landscapes in eukaryotic genomes. Nucleic Acids Research 48, 6685–6698.

Zilberman D, Coleman-Derr D, Ballinger T, Henikoff S. 2008. Histone H2A.Z and DNA methylation are mutually antagonistic chromatin marks. Nature 456, 125–129.

