## Supplementary Figures for "Cas9-targeted Nanopore sequencing rapidly elucidates the transposition preferences and DNA methylation profiles of mobile elements in plants"

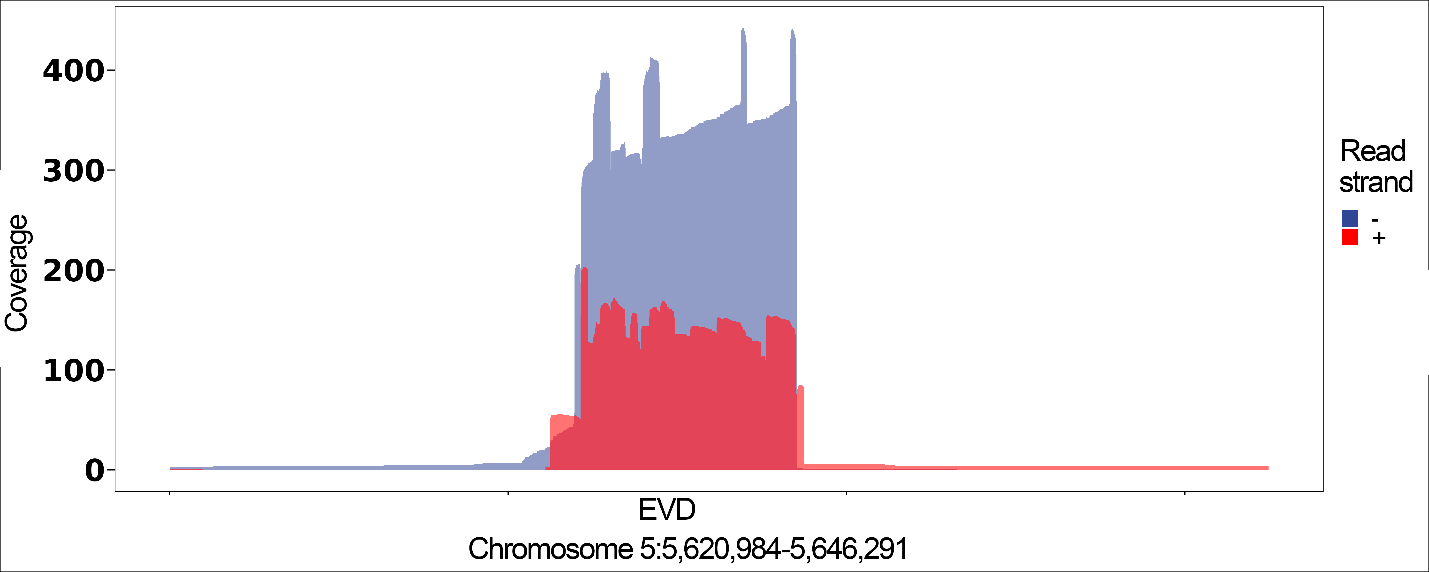

**Supplementary Figure S1.** Coverage plot of EVD5 using forward (red) and reverse (blue) reads obtained following CANS procedure with genomic DNA of G-ddm1-3 plant.

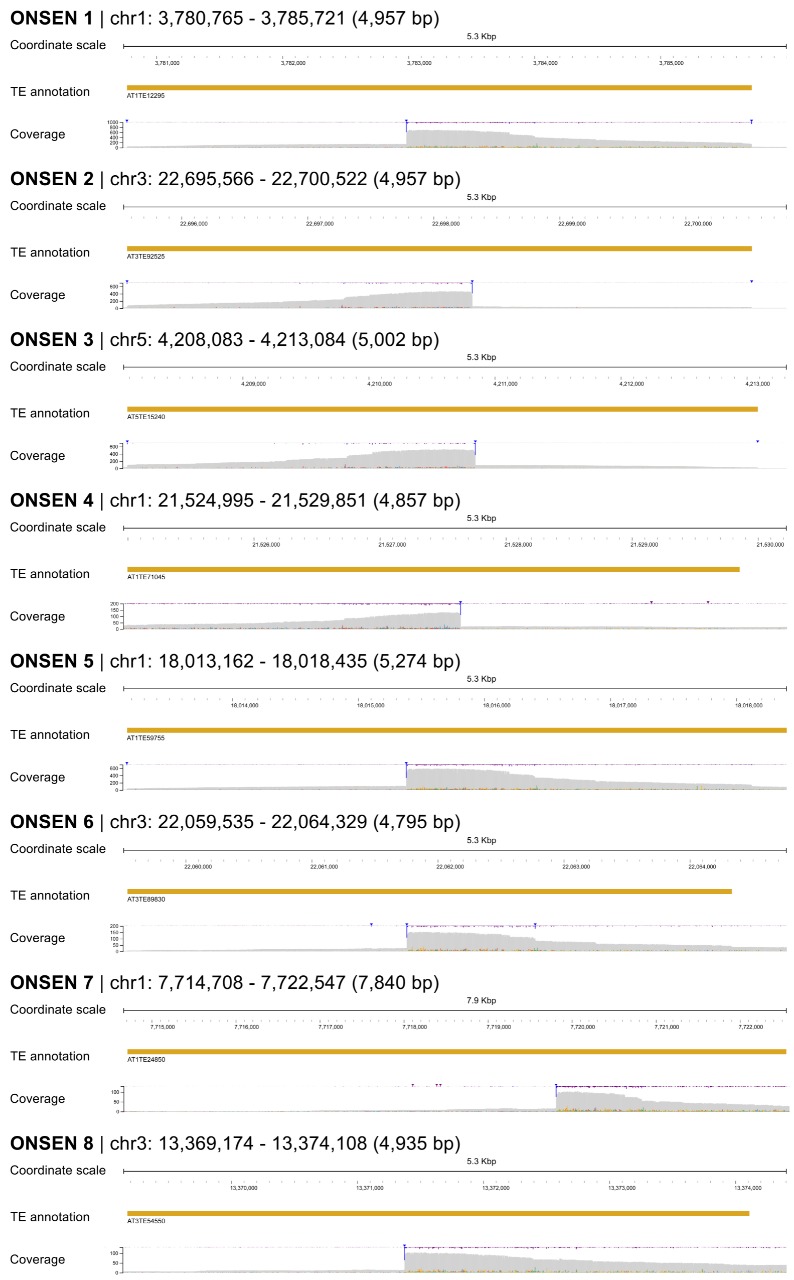

Supplementary Figure S2. Coverage of 8 ONSEN elements by CANS reads.

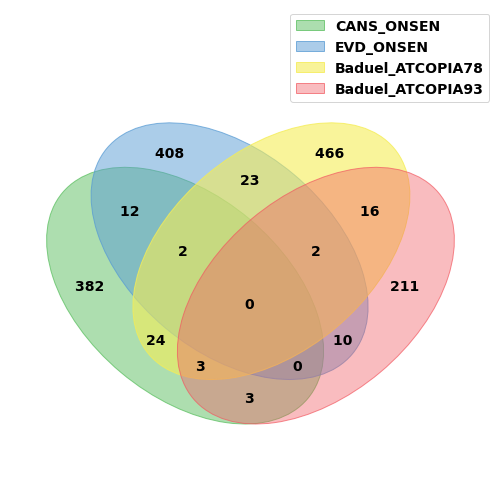

Supplementary Figure S3 . Venn diagram showing number of genes affected by ONSEN and EVD detected by CANS sequencing and short-reads whole-genome sequencing of >1000 Arabidopsis thaliana accessions (Baduel et al. 2021)

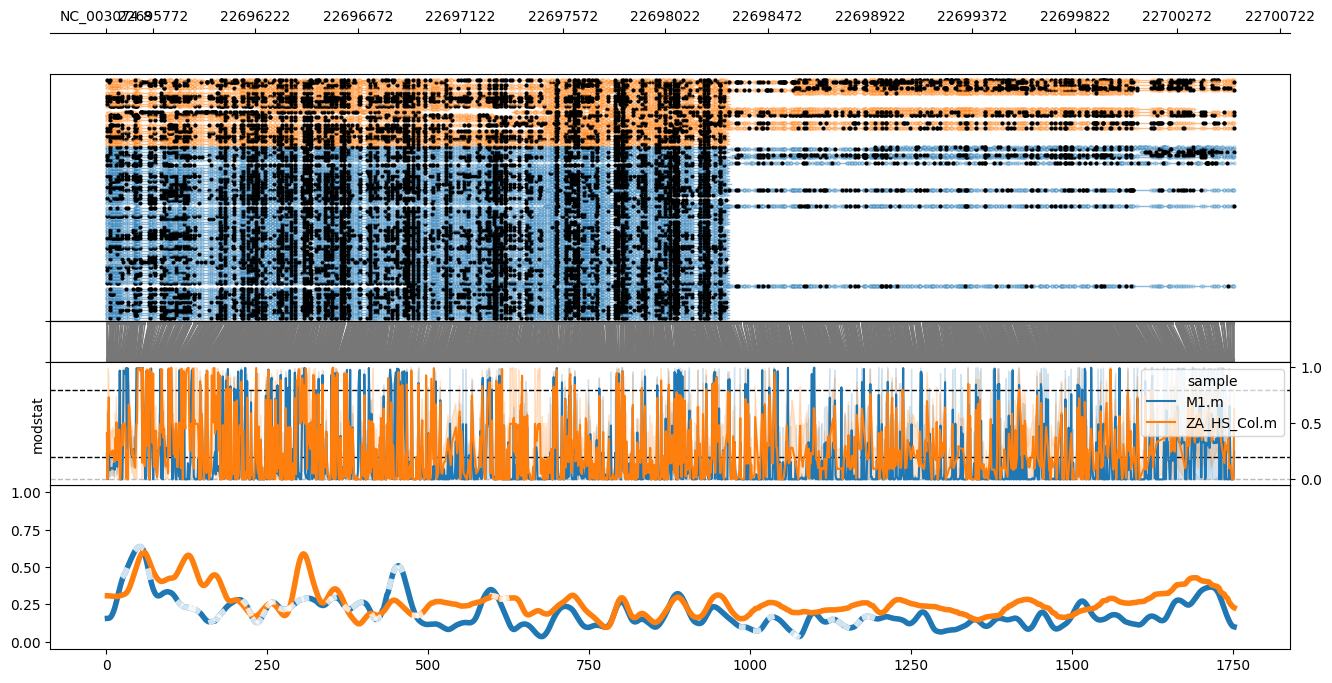

Supplementary Figure S4. CG methylation distribution along ONSEN2 (AT3TE92525, NC_003074.8:22695566-22700522) deduced by M0 (orange) and M1 (blue) CANS reads. From top to bottom: CANS read mapping with indicated methylated (closed circles) and unmethylated (opened circles) cytosine, raw log-likelihood ratios, and smoothed methylated fraction plot.

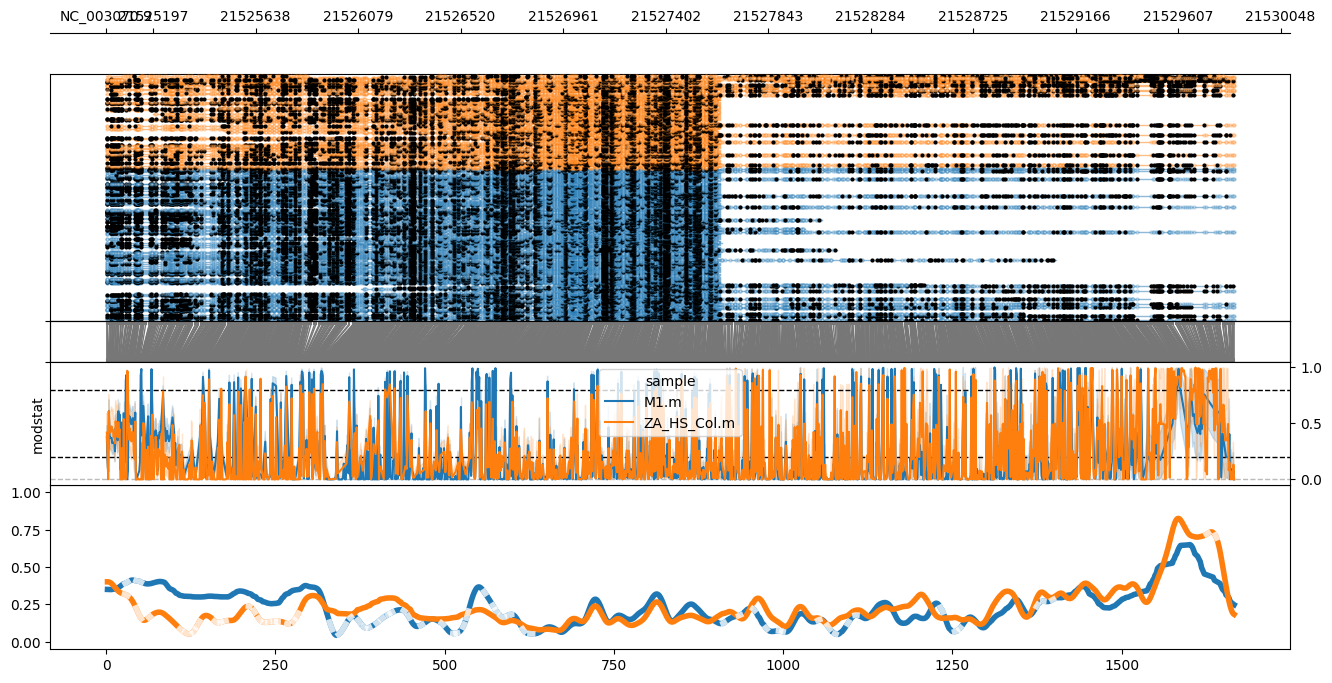

Supplementary Figure S5. CG methylation distribution along ONSEN4 (AT1TE71045, NC_003070.9:21524995-21529851) deduced by M0 (orange) and M1 (blue) CANS reads. From top to bottom: CANS read mapping with indicated methylated (closed circles) and unmethylated (opened circles) cytosine, raw log-likelihood ratios, and smoothed methylated fraction plot.

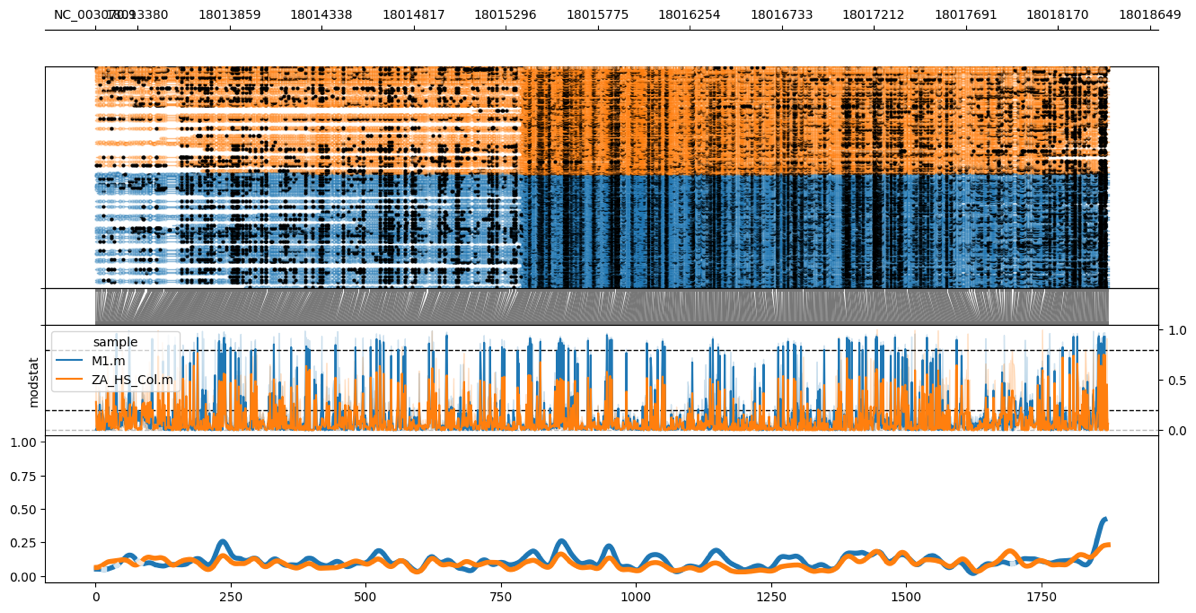

Supplementary Figure S6. CG methylation distribution along ONSEN5 (AT1TE59755, NC_003070.9:18013162-18018435) deduced by M0 (orange) and M1 (blue) CANS reads. From top to bottom: CANS read mapping with indicated methylated (closed circles) and unmethylated (opened circles) cytosine, raw log-likelihood ratios, and smoothed methylated fraction plot.

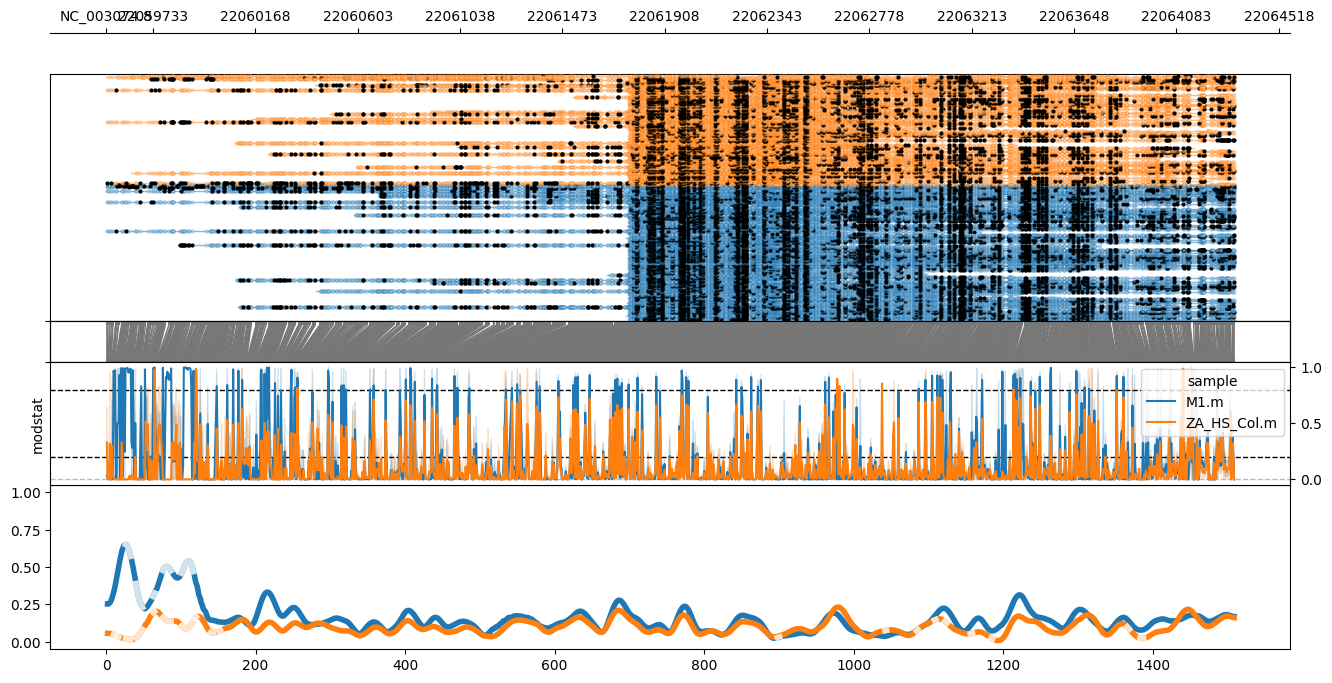

Supplementary Figure S7. CG methylation distribution along ONSEN6 (AT3TE89830, NC_003074.8:22059535-22064329) deduced by M0 (orange) and M1 (blue) CANS reads. From top to bottom: CANS read mapping with indicated methylated (closed circles) and unmethylated (opened circles) cytosine, raw log-likelihood ratios, and smoothed methylated fraction plot.

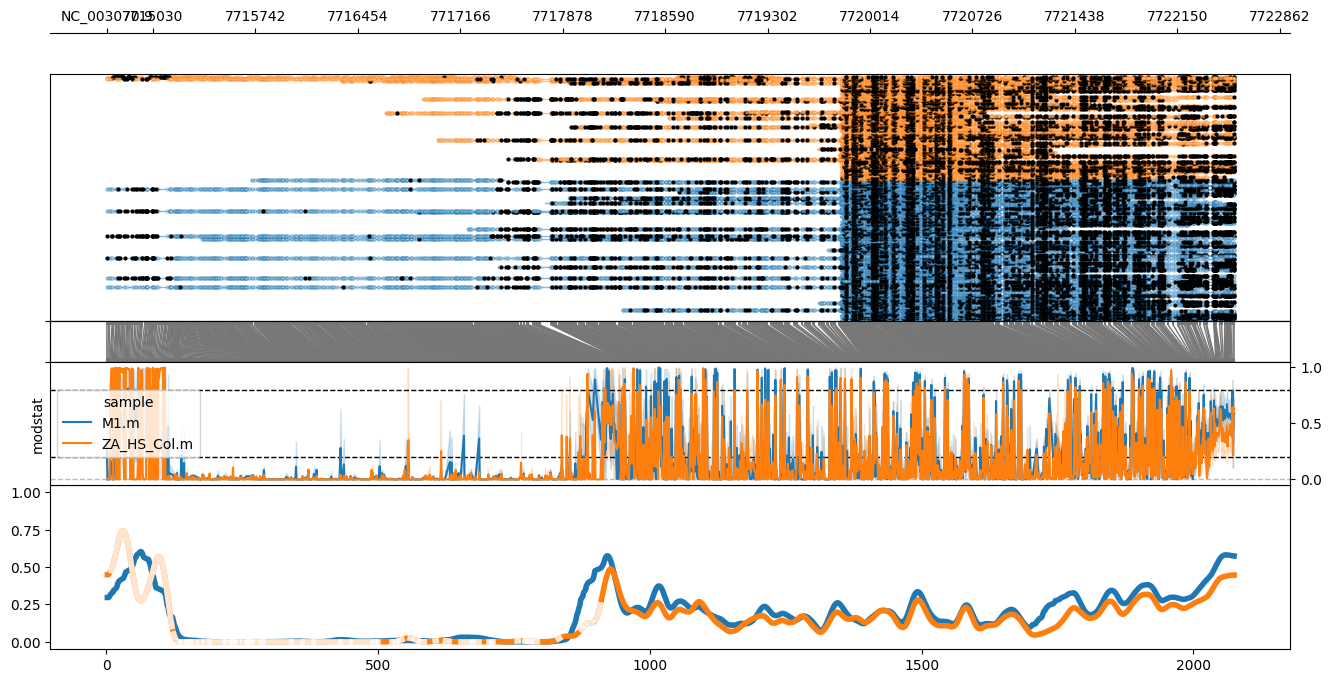

Supplementary Figure S8. CG methylation distribution along ONSEN7 (AT1TE24850, NC_003070.9:7714708-7722547) deduced by M0 (orange) and M1 (blue) CANS reads. From top to bottom: CANS read mapping with indicated methylated (closed circles) and unmethylated (opened circles) cytosine, raw log-likelihood ratios, and smoothed methylated fraction plot. The decrease in CG methylation corresponds to the position of the annotated gene, AT1G21940 as a part ONSEN7 sequence.

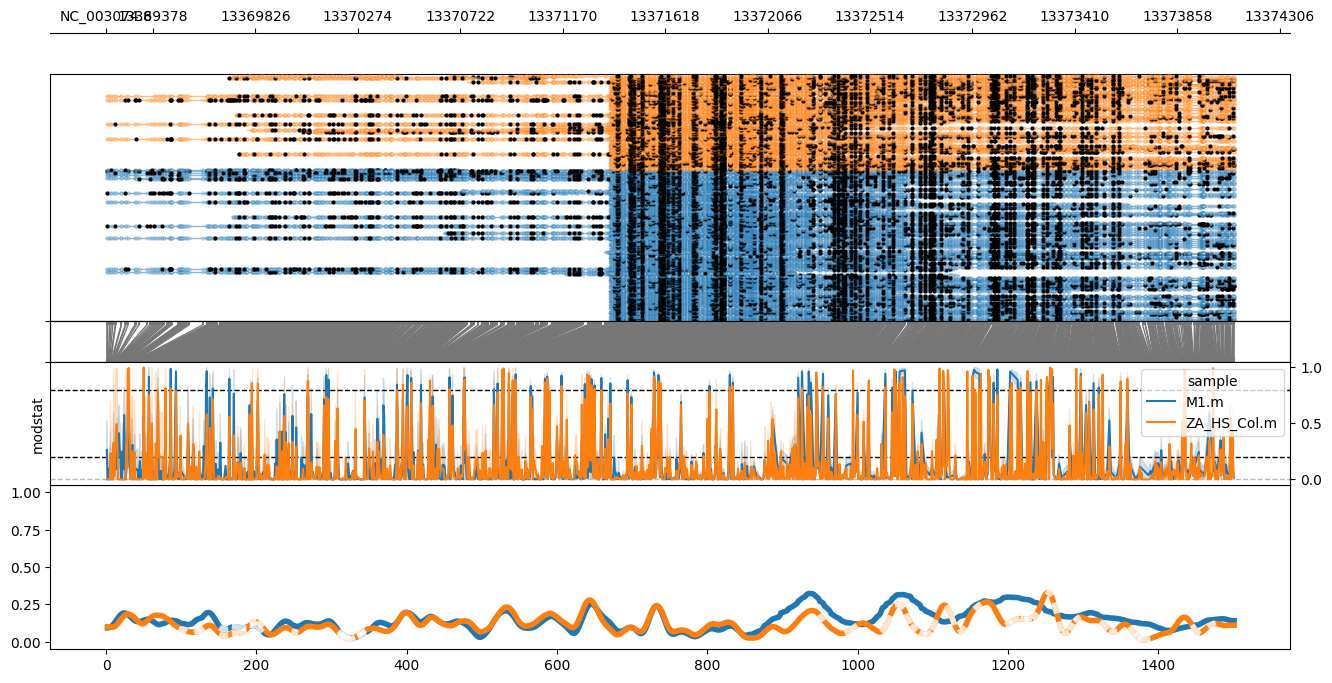

Supplementary Figure S9. CG methylation distribution along ONSEN8 (AT3TE54550, NC_003074.8:13369174-13374108) deduced by M0 (orange) and M1 (blue) CANS reads. From top to bottom: CANS read mapping with indicated methylated (closed circles) and unmethylated (opened circles) cytosine, raw log-likelihood ratios, and smoothed methylated fraction plot.

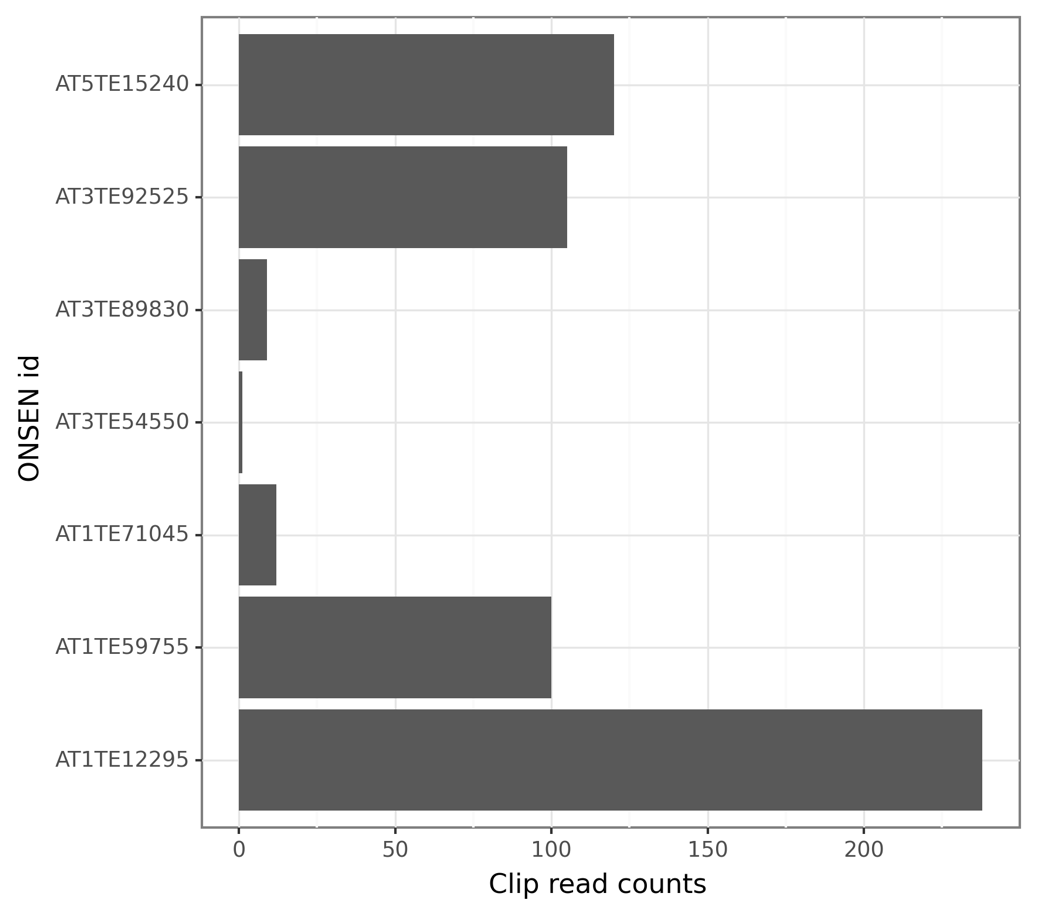

**Supplementary Figure S10. Bar plot showing the number of CANS reads for each ONSEN member corresponding to somatic insertions as determined by NanoCasTE pipeline.**
